# Metabolic reprogramming of protists within the microbiome of sinking particles

**DOI:** 10.1101/2025.05.08.652973

**Authors:** Qingwei Yang, Hisashi Endo, Yanhui Yang, Akiko Ebihara, Yosuke Yamada, Hideki Fukuda, Toshi Nagata, Hiroyuki Ogata

**Affiliations:** Institute for Chemical Research, Kyoto University, Uji, Kyoto, Japan; Atmosphere and Ocean Research Institute, The University of Tokyo, Kashiwa, Chiba, Japan; Institute for Extra-cutting-edge Science and Technology Avant-garde Research, Japan Agency for Marine-Earth Science and Technology, Nankoku, Kochi, Japan

**Author notes:** Corresponding authors: H. Ogata.

**Keywords:** biological carbon pump, metatranscriptome, protists, sinking particles

## Abstract

Protists play a crucial role in the biological carbon pump, yet their metabolic activity and function in sinking particles, which drive the aggregation and degradation of organic carbon, remains poorly characterized. Here, we examined transcriptional dynamics of protists in sinking and suspended particles collected by marine snow catcher in the Oyashio region to explore active lineages and their function during the sinking export. We found taxon-specific metabolic variations, with suspended particles showing up-regulated transcripts from diatoms, haptophytes, and chlorophytes, while sinking particles exhibited those from diatoms, ciliates, and dinoflagellates. Our data further revealed niche-dependent cellular metabolic reprogramming. The small diatoms *Minidiscus variabilis* and *Thalassiosira oceanica* upregulated more genes in suspended particles, while the chain-forming *Thalassiosira rotula* overexpressed more genes in sinking particles. The activity of both phototrophic and heterotrophic protists in sinking particles suggests the export of fresh organic carbon, with heterotrophic protists driving its degradation. Notably, the dinoflagellates *Karlodinium veneficum* and *Karenia brevis* exhibited significantly upregulated genes related to feeding and carbon metabolism in sinking particles. Additionally, the activity of RNA viruses was positively correlated with carbon flux. Overall, our study highlights the critical role of large-celled diatoms, heterotrophic dinoflagellate, ciliates and RNA viruses in organic carbon aggregation and degradation.

## Introduction

As a primary component of the biological carbon pump (BCP) , sinking particles, serving as conduits for the export of photosynthetically produced organic matter, transport carbon and energy to the ocean interior and seafloor, thereby regulating atmospheric CO_2_ level over various time scales [1]. The characteristics of sinking particles depend largely on the composition of phytoplankton community [2] and associated heterotrophic organisms, including zooplankton, prokaryotes, and protists (i.e., unicellular eukaryotes) [3, 4]. These communities influence the efficiency of carbon export by facilitating aggregation, disaggregation, degradation, and trophic transfer [5, 6].

Over the past few decades, the taxonomic composition of marine particle-associated microbes has been extensively analyzed across different particle types with the use of a variety of methodologies [7–14]. A key method in this research field has been the use of large-volume settling chambers, known as Marine Snow Catchers (MSCs), which allow effective separation and analysis of communities on suspended versus sinking particles [8]. Previous studies indicate that marine stramenopiles and oligotrophic prokaryotes prevail in suspended particles, whereas ciliates, dinoflagellates, and copiotrophic prokaryotic generalists are better adapted to sinking particles [12, 14]. Despite these advances, significant knowledge gaps remain with respect to the activity and function of protist community and their impact on the degradation of organic matter in sinking particles.

Previous studies provided insights into the metabolic requirements to take advantage of the microscopic environments in sinking particles especially for bacteria. Particle-associated bacterial communities exhibit significant differences from free-living states, particularly in their enhanced ability to degrade polysaccharides and transport amino acids [15, 16]. Moreover, particles act as hotspots for enzymatic hydrolysis, with hydrolysis rates substantially exceeding those in ambient seawater [5, 17]. However, the extent of metabolic activity exhibited by eukaryotic phytoplankton (diatoms, haptophytes, and chlorophytes) and planktonic predators and heterotrophs such as ciliates and dinoflagellates within sinking particles remains poorly understood. Phytoplankton harness sunlight to convert inorganic carbon into organic compounds. They typically transcribe genes associated with photosynthetic machinery, which fix CO2 and provide the main components of sinking aggregates [18]. Predatory heterotrophs, such as ciliates, are generally motile and consume smaller prey by phagocytosis, thereby enhancing the decomposition of organic matter within sinking particles. This process is characterized by increased expression of genes involved in the degradation and uptake of ingested organic material [19, 20]. A growing number of marine protists, particularly dinoflagellates, are recognized as mixotrophs [21]. These organisms exhibit a range of metabolic strategies that allow them to transform carbon through both photosynthetic fixation in the euphotic zone and decomposition in the mesopelagic layer [21, 22]. However, the diverse trophic modes of protists and their species-specific responses to environmental changes complicate *in situ* characterization of their ecological functions and activities. Metatranscriptomics provides insight into the composition of transcript pools in mixed microbial assemblages [23], allowing the inference of species-specific physiology and the elucidation of links between taxon-level metabolism and ecosystem function.

In this study, we employed a metatranscriptomic approach to analyze the active metabolic pathways of protists within sinking and suspended particles collected by MSCs from Oyashio waters. By comparing *in situ* gene expression profiles of protistan communities in both particles, we identified active metabolic functions, associated these functions to specific protistan species, and assessed the contributions of different protists to the BCP and biogeochemical cycles.

## Method

### Sample Collection

Seawater sampling was conducted during the research cruises KS-21-4 (March 11–21) and KS-21-7 (May 3–11, 2021) by R/V *Shinsei-Maru* (JAMSTEC). A total of 32 samples associated with sinking and suspended particles were collected from four sites with MSCs in the Oyashio region off Hokkaido, Japan (Fig. S1, Table S1). Detailed protocols for field collection and preservation of samples were described in Yang *et al.* (2024). Briefly, sinking particles were collected in the base parts of the MSCs, while suspended particles were collected in the upper parts of the MSCs after standing for 2 h, with sinking speed < 8 m d^-1^ or < 14 m d^-1^. The particles were then filtered through a 0.8 µm membrane and subsequently stored at –80°C. The MSCs were deployed at three depths: the subsurface chlorophyll maximum (SCM, 11–30 m), 10 m below the pycnocline (PYC, 65–250 m), and the bottom boundary layer (BBL, 289–1489 m) (Table S1). Environmental parameters, including particulate organic carbon (POC) and nitrogen (PON) flux were also documented in Yang *et al.* (2024), transparent exopolymer particles (TEP) data were obtained from the same filed study [24].

### Metatranscriptomic sequencing and bioinformatics analysis

Total RNA was extracted using the AllPrep DNA/RNA Mini Kits (Qiagen) and purified with the RNase-Free DNase Set (Qiagen), following the manufacturer’s protocol. The RNA quality was assessed with a BioAnalyzer. Removal of rRNA and isolation of mRNA were performed using a NEBNext® Poly(A) mRNA Magnetic Isolation Module (New England Biolabs, cat. no. E7490) according to the manufacturer’s protocol. Metatranscriptomic libraries were prepared using the NEBNext® Ultra™ II Directional RNA Library Prep Kit for Illumina® (New England Biolabs, cat. no. E7760), according to the provided protocol. Sequencing of all libraries was carried out on the NovaSeq 6000 system (Illumina, USA), generating paired-end reads of 150 bp.

Raw sequences (about 74 million paired-end reads per sample, Table S1) were quality-trimmed using Fastp *(v. 0.23.4, -q 20, -m 50*) [25] and PCR duplicates were removed with FastUniq [26]. Putative rRNA reads were filtered out utilizing SortMeRNA (*v. 4.3.6*) with all recommended rRNA databases [27]. The remaining high-quality reads were *de novo* assembled for each sample by Trinity (*v. 2.15.2*), with a minimum contig length of 300 bp [28]. Assemblies from all samples were combined. Protein-coding sequences for each transcript were predicted across six reading frames using transeq vEMBOSS:6.6.0.059 [29], following the Standard Genetic Code. Only the longest coding sequence from each transcript was used for downstream analyses.

Protein-coding sequences were clustered at a 99% amino acid sequence identity threshold using linclust function of MMseqs2 [30]. From each cluster, the longest protein-coding sequences was retained for further annotation, and its corresponding transcript was selected as a representative transcript. The read counts of transcripts were determined by aligning quality-filtered paired-end reads to representative transcripts with Salmon (*v. 1.10.1*) using the default parameters [31]. Sequences were taxonomically annotated against a custom protein sequence database using DIAMOND (*v. 2.1.9*, *-e 1e-5 -top 10*) [32]. Taxonomic identification and ranking were refined using the lowest-common ancestor in DIAMOND along with NCBI taxonomy. The custom database incorporated MarFERReT [33], NCBI RefSeq [34], RNA virus sequences [35], and large DNA virus sequences from GOEV [36]. For functional annotation, sequences were queried against carbohydrate-active enzymes (CAZymes) and the Kyoto Encyclopedia of Genes and Genomes (KEGG) database [37] using eggNOG [38]. Several KEGG gene categories, including Metabolism (09100), Genetic Information Processing (09120), Environmental Information Processing (09130, excluding the Information Processing in Viruses subcategory), and Cellular Processes (09140), were manually curated for the downstream analysis. Putative biomarker genes for phototrophy and heterotrophy (Table S2) were compiled from the KEGG database and previous studies [39] to examine potential shifts in the trophic strategies of specific protistan taxa during sedimentation. We selected the putative capacity to degrade carbohydrates based on the CAZymes annotation results, including glycoside hydrolases (GHs), carbohydrate esterases (CEs), polysaccharide lyases (PLs), carbohydrate-binding modules (CBMs), and auxiliary activities (AAs).

### Protistan community analysis

Transcript expression levels were normalized by two approaches: transcripts per million (TPM), accounting for transcript length and sequencing depth to facilitate comparison of transcript abundance across samples, and the trimmed mean of M values (TMM) method [40], adjusting for library size and compositional variances to ensure precise comparisons of gene expression across different samples. The similarity of metatranscriptome communities was assessed using non-metric multidimensional scaling (nMDS) of Bray-Curtis dissimilarity, applying TPM-normalized read counts through the vegan package [41], with metazoan transcripts being excluded to focus on protists. The statistical significance of differences between sinking and suspended particles was evaluated using permutational multivariate analysis of variance (PERMANOVA) with 9999 permutations. Differentially expressed gene (DEG) analysis between particle types was conducted using edgeR [42], applied to a TMM-normalized dataset from which low-expressed transcripts were removed. Transcripts with a false discovery rate (FDR) adjusted *p*-value < 0.05, as calculated by the Benjamini-Hochberg procedure [43], and a minimum fold change (FC) of >1.5 were identified as significantly differentially expressed.

### Taxonomic bins analysis

Eukaryotic transcripts from the taxonomy assignment were further aligned to MarFERReT [33] using DIAMOND blastp [32] to recover species-level taxonomic bins. We retained the best hits for each transcript with >95% identity. Taxonomic bins with low activity were filtered based on having >1,000 non-zero transcripts with KO hits in ≥12 samples (6 paired sinking vs. suspended). Low-abundance samples and their paired counterparts were excluded. This process resulted in 26 species-level taxonomic bins, which were used for subsequent DEG analysis, following the same method as the community-level analysis.

### Detecting Protistan viruses

The viral and unclassifiable transcripts identified in the taxonomy assignment step were selected for further detecting signals of giant viruses, single-strand DNA (ssDNA) viruses, and RNA viruses, which are considered the primary viruses infecting protists. Protein-coding sequences were re-predicted using prodigal-gv (*v. 2.11.0*) [44] to improve gene calling for viruses. Viral marker and core genes were identified by searching against HMM profiles (Table S3) with HMMER (*v. 3.4,* http://hmmer.org). The newly detected viral genes, along with taxonomy-assigned viral sequences, were considered viral transcripts.

### Availability of data

All raw sequences are publicly available at the DDBJ Sequence Read Archive under accession number DRA020463.

## Results and Discussion

### Metatranscriptome profiling and taxonomic composition of transcripts

To investigate the metabolic activity of protist community associated with BCP, eukaryotic metatranscriptomic samples were collected from sinking and suspended particles in the Oyashio waters using MSCs during March and May of 2021. In total, we yielded over 1,166 million quality-filtered paired-end reads (Table S1). The assembly of these reads generated 47.0 million nucleotide contigs across samples. Subsequently, the translated sequences were clustered at 99% amino acid sequence identity, resulting in a pool of 16,237,500 representative sequences. Approximately 62.0% of the transcripts were identified as eukaryotic through DIAMOND searches, with the remainder classified as viruses (0.1%), bacteria (0.2%), or unclassifiable (37.8%) (Fig. S2a). The viral and unclassifiable transcripts were further analyzed to refine and augment viral signal detection using HMM searches, ultimately identifying 0.18% of the transcripts as viral. Within the eukaryotic group, approximately 18.4% were assigned to metazoans (Fig. S2b), and about 19.3% had ambiguous taxonomy, which were only annotated at “Eukaryote” level. We focused on the protists (∼60%) by excluding all metazoan and ambiguously annotated transcripts from subsequent analyses. Protistan transcripts were annotated using KEGG orthology (KO), with 29.8% of those being assigned to KOs (Fig. S2c).

Metabolically active protistan groups in Oyashio waters included Bacillariophyceae (diatoms), Dinophyceae (dinoflagellates), Haptophyta (haptophytes), Ciliophora (ciliates), and Chlorophyta (chlorophytes) (Fig. 1). Among these, diatoms were the most abundant in the protistan transcript pools at both Stations 3 and 4 (St3 and St4) in March, representing 50.5%–81.8% of the transcripts in both particle types across depths. This finding is consistent with the occurrence of a diatom bloom at St4 and St3 in March, as indicated by the previous 18S metabarcoding analysis and chlorophyll *a* concentration [14]. The dominance of diatoms’ transcripts in both sinking and suspended particles further supports their central role in the downward flux of organic material during the spring bloom. In other samples (St1 in March and all the stations in May), various protist groups contributed to the transcript pools, with haptophytes averaging 15.4%, dinoflagellates 36.6%, and diatoms 22.7%. Notably, ciliate transcripts comprised around 2.5% of the protistan transcripts, which was much lower than the proportion found in the 18S metabarcoding analysis (17.6%) [14]. This discrepancy may be due to the limited availability of reference gene data for ciliates [33].

**Figure 1.**
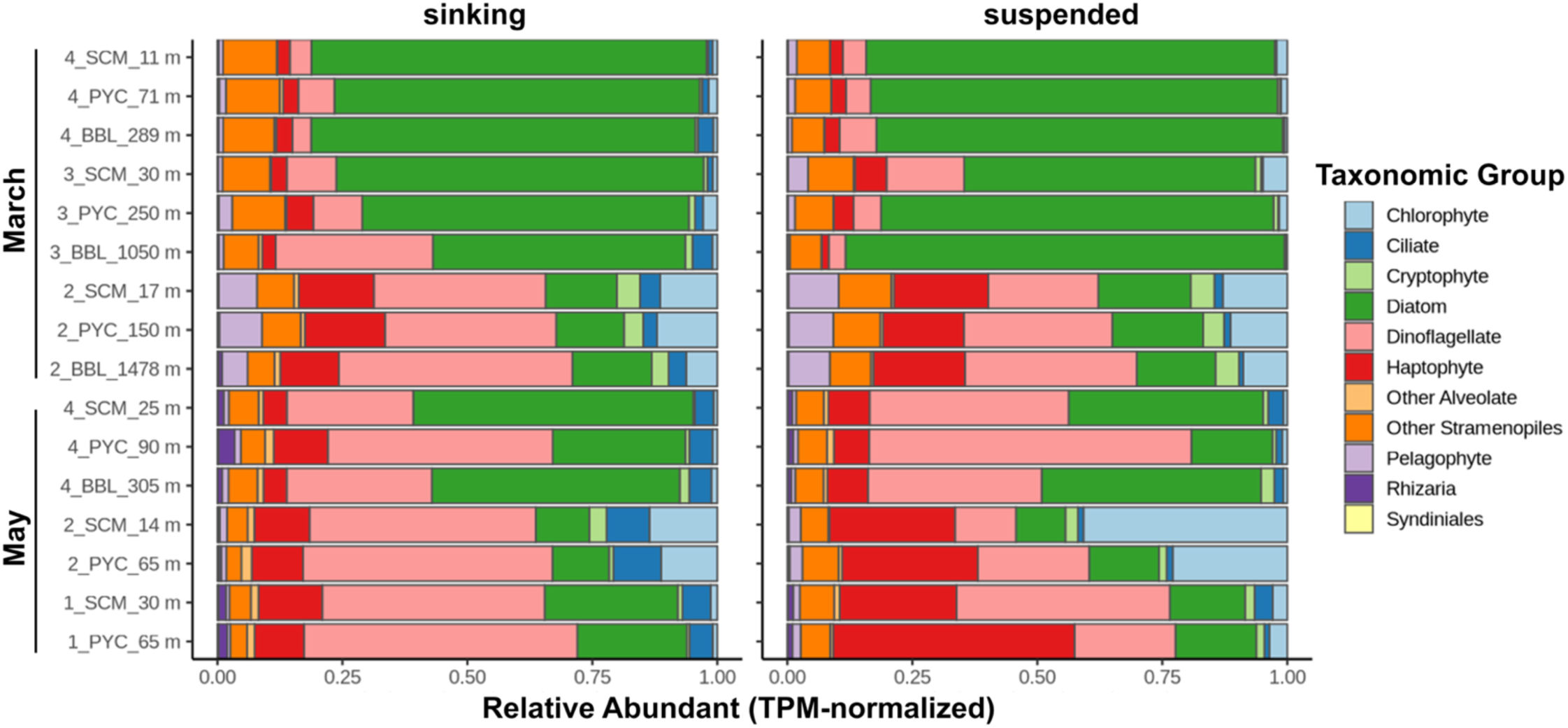
Transcript abundance of major protist taxa in sinking and suspended particles. The relative transcript abundance for each taxon was calculated as the sum of transcripts per million (TPM) annotated to that taxon, divided by the total TPM across all eukaryotes (excluding metazoan and ambiguous eukaryotic transcripts). The x-axis displays the sample information, e.g., “4_SCM_21 m” indicates St4 at the SCM layer, with a depth of 21 m.

### Differential expression pattern of protistan community between particle types

An nMDS analysis was performed overall protistan transcript pools to examine the community-level expression pattern of protists across depths and particle types. The results revealed a significant difference in protistan transcript composition between sinking and suspended particles (PERMANOVA, P < 0.05; Fig. 2a, Table S4), with no obvious significant difference across depths (PERMANOVA, P=0.91; Fig. 2a, Table S4). This distinction of particle types was further confirmed by differential expression analysis. In total, 24,612 transcripts (15.4% of all the transcripts analyzed) were identified as DEGs, of which 9,622 contained KO hits (Fig. 2b and Fig. S3). Among those with KO hits, 7,251 transcripts were significantly more expressed in sinking particles, whereas 2,371 were significantly more expressed in suspended particles (FDR < 0.05 and |logFC| > 1.5, Fig. 2b and Fig. S3). These results highlight the functional differences in protist communities between sinking and suspended particles. Notably, overexpressed transcripts in sinking particles were predominantly associated with diatoms (45.3%), ciliates (24.8%), and dinoflagellates (16.8%), while highly expressed transcripts in suspended particles were mainly linked to diatoms (49.4%), haptophytes (23.8%), and chlorophytes (10.3%) (Fig. 2c-g and Fig. S3). The DEG analysis, conducted specifically for SCM layer (Fig. S4a) and the combined PYC and BBL layers (Fig. S4b), provided additional evidence supporting these taxon-specific expression patterns. These notable taxon-specific expression patterns between particle types further support the selective shaping of the protistan community in the aggregate biosphere as indicated by 18S metabarcoding analysis from the same samples [14].

**Figure 2.**
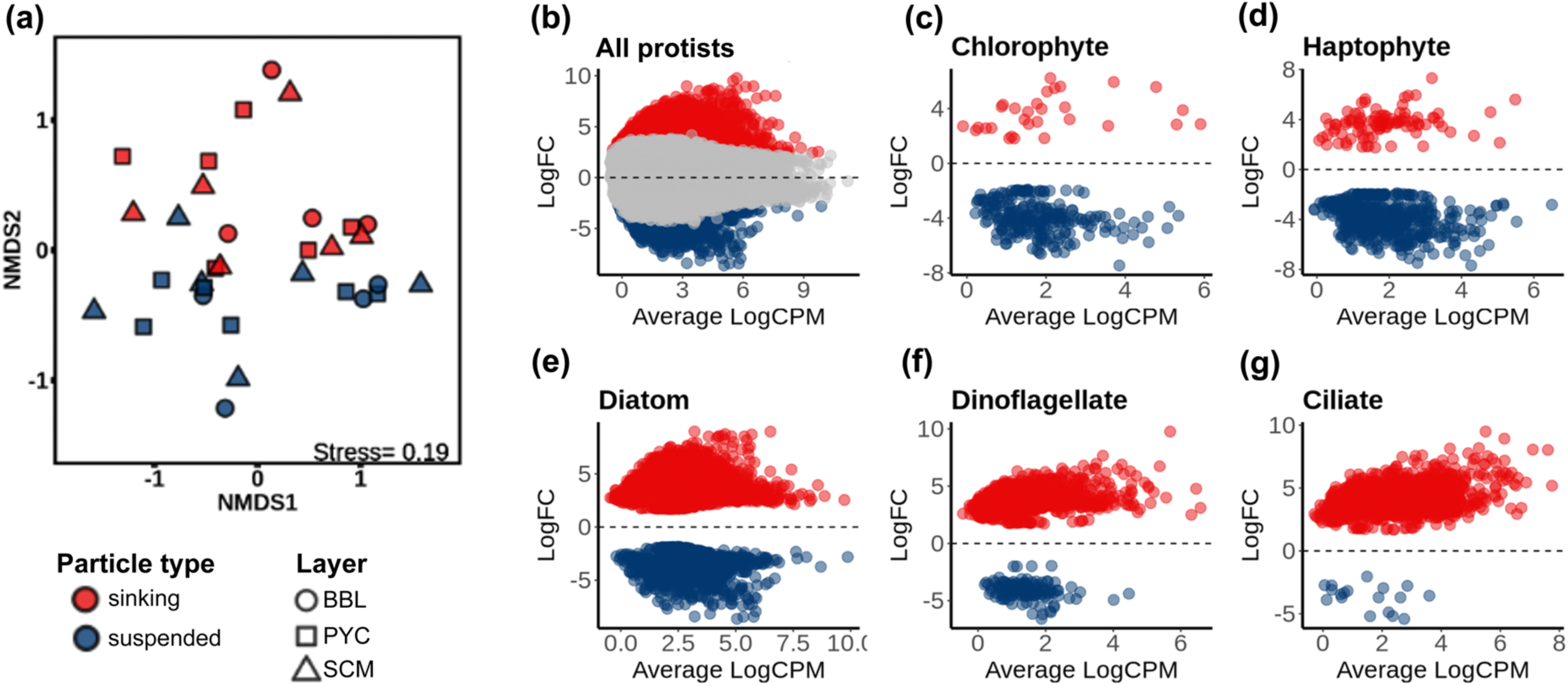
Differential expression between sinking and suspended particles. (A) nMDS plot showing differences in protistan community composition between sinking and suspended particles, and across depths, based on transcript expression levels (TPM-normalized). (B) MA plot illustrating log2 fold changes (logFC, y-axis) between sinking and suspended particles for all protistan transcripts, plotted against the mean of log-transformed counts per million (CPM) from TMM-normalized libraries (only KO-annotated transcripts included). Colored markers indicate differentially expressed transcripts (FDR < 0.05 and |logFC| > 1.5), with higher expression in sinking (red) and suspended (blue) particles. (C-G) The MA plot from (b) showing differentially expressed transcripts, further divided by taxa: (C) Chlorophytes, (D) Haptophytes, (E) Diatoms, (F) Dinoflagellates, and (G) Ciliates

A detailed functional analysis of DEGs in sinking particles revealed a high proportion of up-regulated DEGs from dinoflagellate and ciliate associated with the function of transport and catabolism (e.g., lysosome and phagosome pathways), cell growth and death (e.g., apoptosis), and signal transduction (e.g., MAPK, PI3K-Akt, and AMPK signaling pathways) (Fig. 3). Similarly, in suspended particles, the function of up-regulated DEGs from haptophytes and chlorophytes were associated with cell growth and death (e.g., apoptosis), transport and catabolism (e.g., lysosome and phagosome pathways), and signal transduction (Fig. 3). These findings highlight potential mechanisms underlying the selective aggregation and disaggregation of sinking particles associated with various protists and their functions.

**Figure 3.**
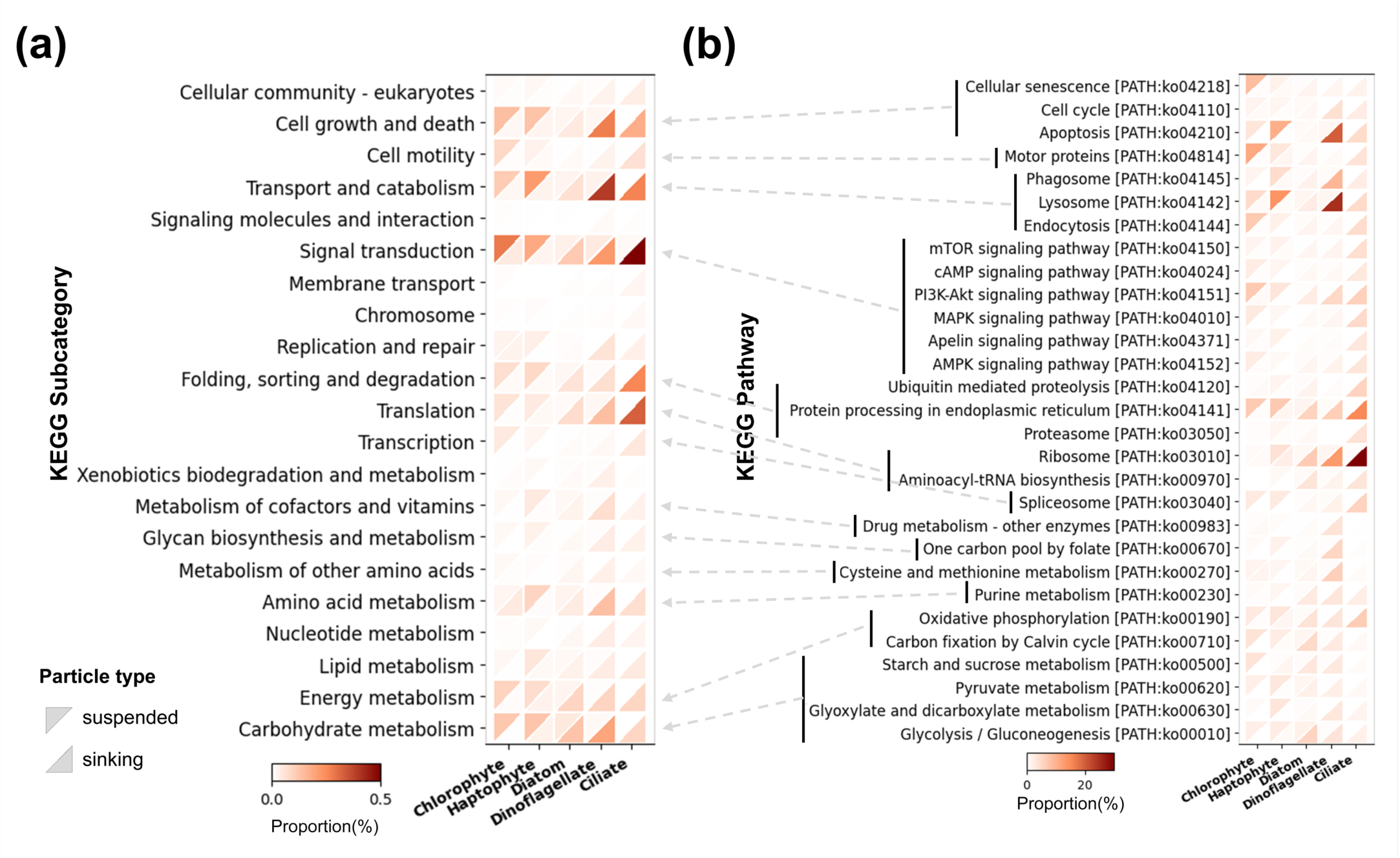
Heatmap showing the functional distribution of differentially expressed genes from Figure 2b within each major taxon. (A) Functions were grouped by KEGG subcategories. (B) Functions were further grouped by KEGG pathways, retaining only those pathways that accounted for more than 3% of the total in a single taxon and contained more than 20 KO counts in that taxon. The dashed arrows indicate the correspondence between KEGG pathways and their respective KEGG subcategories.

Firstly, size-dependent predator-prey interactions [45, 46] may influence predators’ preference for sinking versus suspended particles. Phagosomes are membrane-bound vesicles that form around a particle during the process of phagocytosis [47]. Lysosomes are membrane-bound cytoplasmic organelles containing hydrolytic enzymes that are responsible for the degradation of various biomolecules [48]. For dinoflagellates and ciliates, we observed an increased proportion of up-regulated DEGs related to phagosome and lysosome pathways, as well as signal transduction in sinking particles (Fig. 3). However, autophagy-related DEGs, associated with starvation in protists [49], were barely detected. These results suggest that dinoflagellates and ciliates in sinking particles are not starving and exhibit more pronounced predatory behavior compared to suspended particles. This observation is consistent with earlier studies, which documented that heterotrophic protists preferentially aggregate on marine snow to enhance feeding [50]. Additionally, we observed a relatively high proportion of up-regulated DEGs associated with function of translation (e.g., ribosomes) in ciliates from sinking particles (Fig. 3). As previous study has shown a linear correlation between ribosome abundance and growth rate in several organisms [51], our result suggest that ciliates had a relatively high growth rate in the sinking particles. Photosynthetic picoeukaryotes, such as chlorophytes and haptophytes, show relatively high expression of phagosome, lysosome, and motor proteins in suspended particles (Fig. 3). These picoeukaryotes have been documented to be capable of phago-mixotrophy, enabling them to feed on small particles like bacteria [52–54]. This suggests that some of the species of chlorophytes and haptophytes in our samples may adopt a mixotrophic lifestyle, favoring a heterotrophic or grazing strategy in a free-living (“suspended”) state. Previous studies indicate that for bacterivorous protists, feeding on bacteria attached to aggregates is more energetically demanding than grazing on freely suspended bacteria, as a relevant additional effort is required [55].

Secondly, the programmed cell death (PCD)-mediated mortality in protists may promote aggregation, with the sinking velocity of these aggregates being influenced by plankton composition. A previous study has shown that PCD in cyanobacteria leads to a significant increase in the excretion of transparent exopolymers, contributing to substantial carbon flux to deeper ocean layers [56, 57]. In haptophytes and dinoflagellates, the elevated expression of genes associated with apoptosis was observed in suspended and sinking particles, respectively (Fig. 3). These apoptosis-associated genes may actually refer to process relate to PCD in protists. Thus, we speculate that PCD-driven mortality of protists also facilitates their aggregation. However, due to the smaller cell size of haptophytes and the lower density of transparent exopolymer, which may enhance particle suspension [24], haptophyte-enriched aggregates likely exhibit lower sinking velocities. Consequently, these aggregates are more prone to being collected as suspended particles (with sinking velocities < 8 m d^-1^ and < 14 m d^-1^) in our samples.

### Vertical dynamics of phototrophy- and heterotrophy-associated transcripts in protistan communities

We analyzed the vertical dynamics of trophic strategies in protists, based on phototrophy- and heterotrophy-related gene expression in sinking and suspended particles, and investigated of the contributions of the lifestyles to the degradation of organic carbon in the sinking particles during sedimentation. The expression of phototrophy- and heterotrophy-related transcripts constituted 16.8% ± 2.8% of the protistan transcript pools. The diatom, haptophyte, and chlorophyte groups comparably expressed both phototrophy- and heterotrophy-related genes, whereas dinoflagellates and ciliates primarily expressed genes associated with heterotrophy (Fig. 4). Notably, many phototrophy-related genes are plastid-encoded and not enriched by poly-A-selected sequencing, potentially leading to an underestimation of phototrophy-related genes expression.

**Figure 4.**
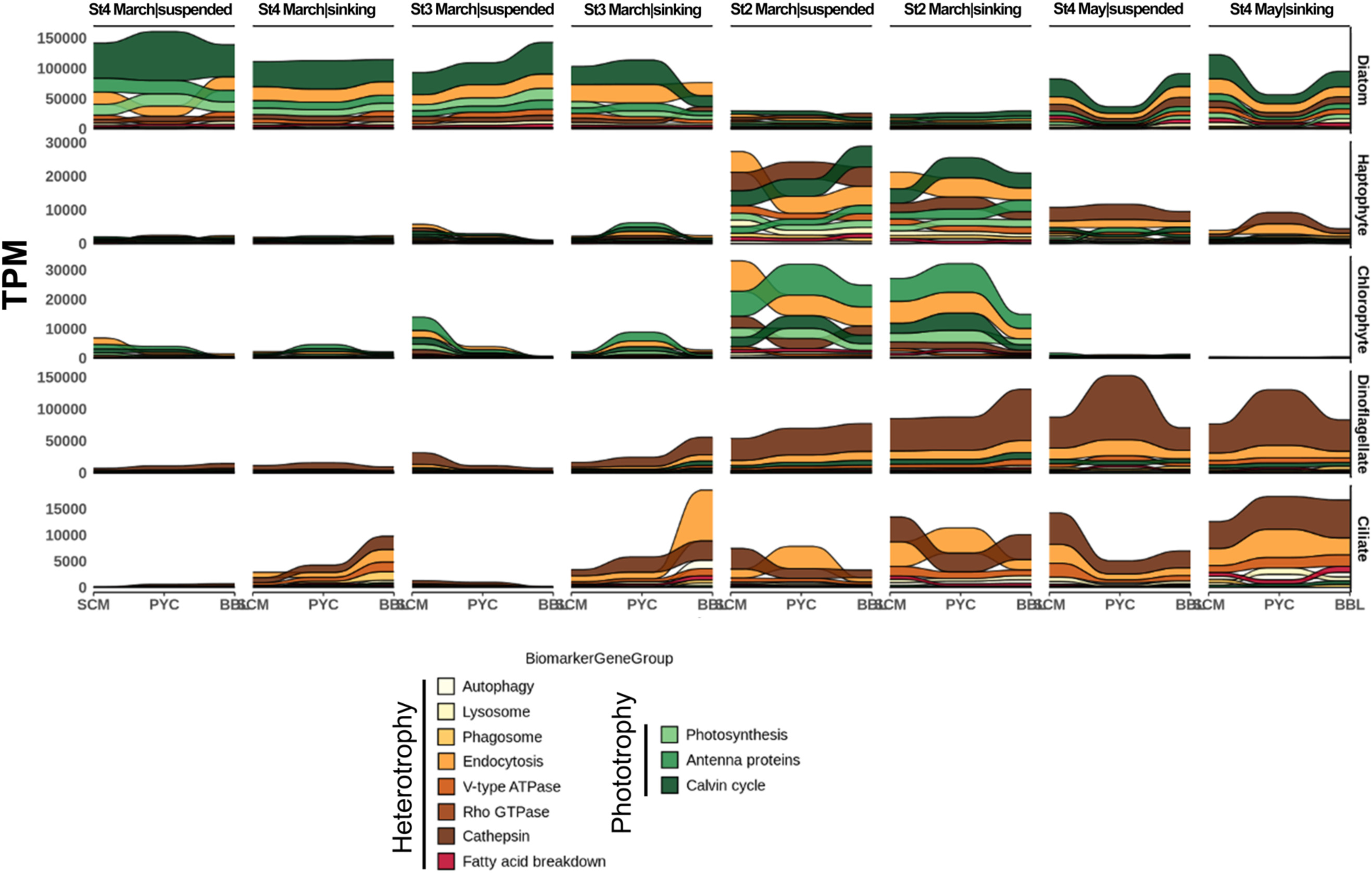
Vertical changes in the expression of various phototrophy and heterotrophy biomarker gene groups across major taxa in sinking and suspended particles. Transcript expression was normalized to TPM. The KEGG KO terms associated with each gene group are provided in Table S2. Stations without a BBL sample (St1 and St2 in May) were excluded from the analysis.

Specifically, the relative expressions of phototrophy-related genes from diatoms, including those associated with photosynthesis, antenna proteins, and the Calvin cycle, were high in both sinking and suspended particles across the water column, down to depths greater than 1000 m (Fig. 4). This result suggests that sinking particles deliver fresh organic carbon, including significant quantities of active phytoplankton, to deeper ocean layers, where they likely disaggregate. In agreement with these findings, Agusti *et al.,* (2015) documented the widespread presence of healthy photosynthetic cells, primarily diatoms, at depths of down to 4,000 m in the deep ocean. Furthermore, among these potential photosynthetic taxa, an elevated expression of genes associated with heterotrophic marker genes, such as cathepsin and endocytosis, was observed in both sinking and suspended particles across the water column (Fig. 4). Cathepsins are key lysosomal proteases that are primarily located in acidic endo/lysosomal compartments. They play a vital role in intracellular protein degradation and energy metabolism [59, 60]. Endocytosis is a process occurring at the cell surface that allows for the internalization of the plasma membrane along with its associated membrane proteins and lipids [61, 62]. This process is crucial for sampling the extracellular environment and regulating various cellular functions initiated at the cell surface. Diatoms, which are typically regarded as exclusively phototrophic, have been demonstrated to employ endocytosis for the uptake of siderophore-bound iron [63, 64]. It is also conceivable that this iron uptake mechanism may be employed by other eukaryotic phytoplankton, such as haptophytes and chlorophytes [65]. Moreover, as previously mentioned, certain species of haptophytes and chlorophytes demonstrate phago-mixotrophic capabilities, enabling them to utilize endocytosis to capture and ingest small prey, such as bacteria, from sinking particles. In addition, transcripts involved in amino acid metabolism and carbohydrate metabolism (e.g., glycolysis, gluconeogenesis, and citrate cycle) were highly expressed in both sinking and suspended particles throughout the water column (Fig. S5-S6), accompanied by an increased expression of genes associated with GHs (Fig. S7). These findings suggest that the maintenance of essential cellular processes in these potential photosynthetic taxa play a pivotal role in the breakdown of organic carbon within sinking particles.

Although mixotrophic nutrient strategies are common in dinoflagellates, and some ciliates possess mixotrophic capabilities through algal endosymbionts [21, 66], most dinoflagellates and ciliates in our samples appear predominantly heterotrophic. Their relative gene expression levels were significantly higher in sinking particles compared to suspended particles (Fig. 2f-g and 4), indicating greater metabolic activity in sinking particles. We observed the expression of transcripts related to cathepsins, endocytosis, and V-type ATPase increased with depth in the water column and was consistently higher in sinking particles than that in suspended particles (Fig. 4). Similarly, the relative expressions of genes associated with energy metabolism, amino acid metabolism, lipid metabolism, and carbohydrate metabolism (e.g., glycolysis, gluconeogenesis, and citrate cycle) were also elevated in sinking particles and increased with depth, along with an increase of transcripts related to GHs and CEs (Fig. S5-7). These findings suggest that dinoflagellates and ciliates maintain higher energy acquisition and consumption. They may actively graze the photosynthesis cells like diatom and haptophytes [67–70], serving as primary protistan consumers of organic matter during sedimentation. This further highlighted their important role in the degradation and cycling of organic carbon in the BCP, as suggested by previous study [14].

We examined shifts in phototrophy- and heterotrophy-related gene expression through DEG analysis to assess potential adaptation of protists as they sink. However, no DEGs were found between SCM and BBL in either sinking or suspended particles (Fig. S8). This lack of change may be due to limitations, including a small sample size (four paired samples from different stations), which reduced statistical power, and variability in community composition, which may have obscured trends.

### Species-specific expression patterns in sinking and suspended particles

To further investigate the role of protists in the BCP, we analyzed the differential gene expression between sinking and suspended particles at the species-level. In this way, we attempted to disentangle the effects of community shifts and transcriptional reprogramming on the overall trend in the metatranscriptomic landscape at the community level. Of 26 recovered species-level taxonomic bins, eleven were assigned to diatoms, four to dinoflagellates, and the remaining to haptophytes (2), chlorophytes (4), dictyochophytes (3), pelagophytes (1), and bolidophytes (1) (Fig. S9).

Diatom species exhibited varying patterns of differential metabolic activities between sinking and suspended particles (Fig. 5 and Fig. S9). Small centric diatoms, such as *Minidiscus variabilis* (diameter < 5 µm [71]) and *Thalassiosira oceanica* (diameter: ca. 5 µm [72]), showed more up-regulated DEGs in suspended particles, while *Thalassiosira rotula*—a typically chain-forming diatom [73]—exhibited more active genes in sinking particles (Fig. 5). Other diatom species, like *Odontella aurita* and *Detonula confervacea*, showed minimal differences in their transcriptomic profiles between the two particle types (Fig. S9). These results aligns with a previous simulation based hypothesis that the contribution of diatoms to carbon export varies among species [74]. Factors such as cell size, shell thickness, Si/C ratio [74], iron availability, nutrient conditions [75], chain formation [76], and viral infections [77] may all influence such a variation in transcriptional reprogramming across species. We identified DEGs associated with various pathways, including signaling pathways and tight junctions, in these three diatom species (Fig. S10). However, the specific mechanisms behind these differential expressions pattern in diatoms remain unclear and require further investigation.

**Figure 5.**
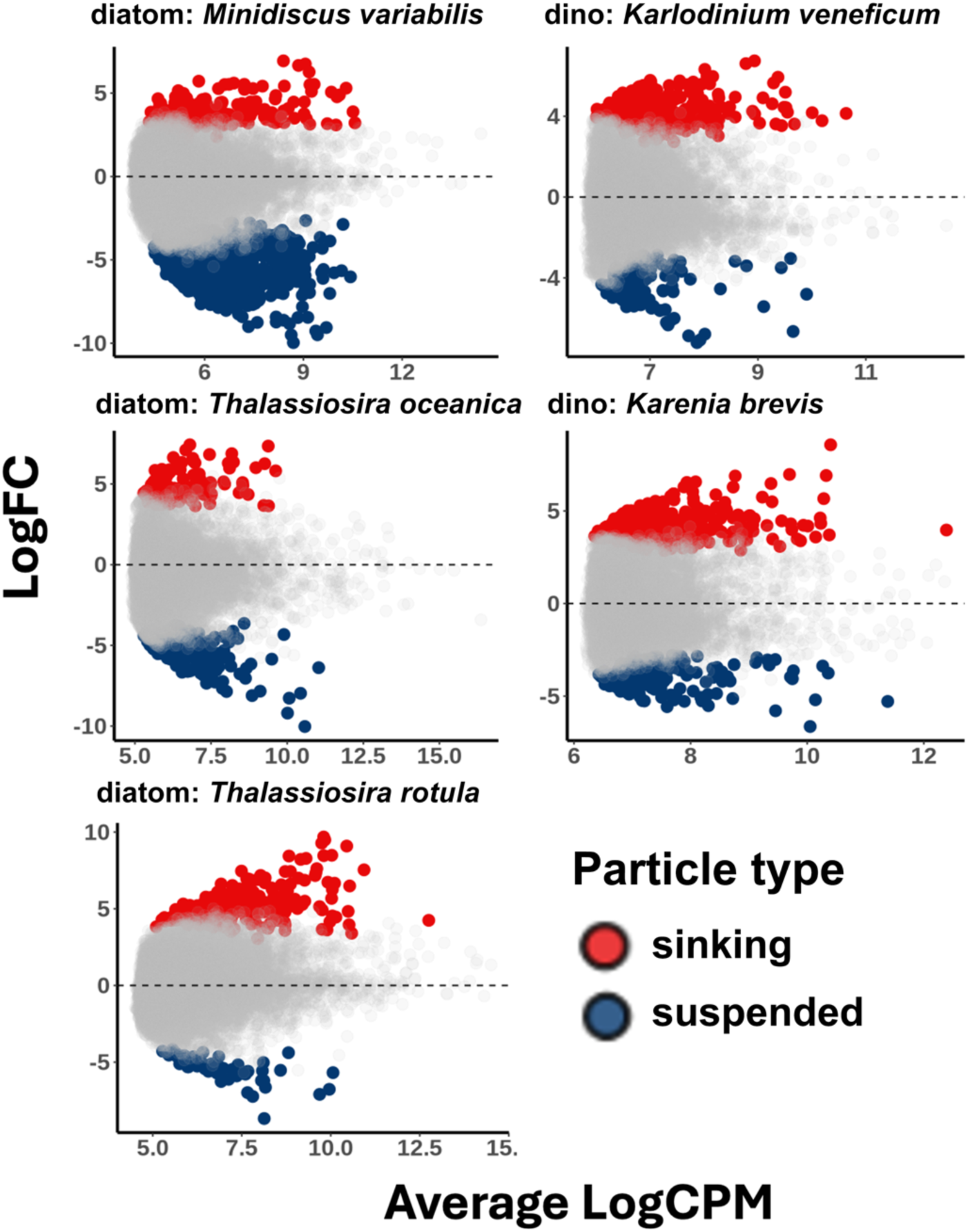
the MA plots show the differential expression of species-specific transcript bins between sinking and suspended particles. The y-axis represents log2 fold change (logFC), and the x-axis shows the mean of log-transformed counts per million (CPM) from TMM-normalized libraries. Only KO-annotated transcripts are included. Colored markers indicate differentially expressed transcripts (FDR < 0.05 and |logFC| > 1.5), with higher expression in sinking (red) and suspended (blue) particles.

Of the four dinoflagellate transcript bins, *Karlodinium veneficum* and *Karenia brevis* showed many up-regulated DEGs in sinking particles (Fig. 5 and Fig. S9). We then examined the function of differentially expressed transcripts for *K. veneficum* and *K. brevis* to determine whether these dinoflagellates species exhibit higher expression of transcripts associated with feeding and the degradation of organic carbon in sinking particles. We found that *K. veneficum and K. brevis* exhibited higher expression of genes related to motor protein, cytoskeleton in muscle cells, the signaling pathways (e.g., MAPK, AMPK, and PI3K-Akt), phagosome, and endocytosis in sinking particles (Fig. S11). The MAPK signaling pathway has been shown to regulate ciliary beat in various eukaryotes [78] and the AMPK signal pathways is a sensor of cellular energy and nutrient status [79]. Both dinoflagellate species have known mixotrophic capacities [80, 81], allowing them to feed on primarily phototrophic organisms, such as cryptophytes. Based on these findings, we speculate that *K. veneficum* and *K. brevis* increases ciliary movement to approach phytoplankton-rich aggregates, thereby feeding on the phytoplankton attached to particles. Additionally, we observed up-regulated DEGs involved in carbohydrate metabolism (e.g., glycolysis, gluconeogenesis, and citrate cycle) in sinking particles (Fig. S11), suggesting that *K. veneficum* and *K. brevis* actively consume organic carbon within sinking particles.

The transcript bins corresponding to haptophytes, chlorophytes, and pelagophytes showed no significant difference between sinking and suspended particles (Fig. S9), although community-level analysis revealed these taxa to be more active in suspended particles (Fig. 2c-d). These inconsistencies may be due to limitations in the metatranscriptomic method for recovering species-level taxonomic bins for DEGs analysis, which included only 11.20% of transcripts.

### Viral activity in sinking and suspended particles

The nMDS analysis of viral transcript composition revealed significant differences in viral activity between sinking and suspended particles (PERMANOVA, P < 0.05, Fig. 6a). This indicates that viruses may play a role in particle aggregation and potentially influence the selective structuring of protistan communities within sinking particles. Among the viral transcripts, the RNA-dependent RNA polymerase (RdRp) was the most highly expressed functional genes (Fig. S12), suggesting that RNA viruses are the most active viral group in the Oyashio waters. We calculated the correlation between viral activity markers—major capsid proteins (MCP) of nucleocytoviruses, HK97 MCP of mirusviruses [36], replication-associated protein (Rep) of ssDNA viruses, and RdRp—and POC/PON flux, as well as TEP, using Pearson correlation. TEP showed a positive correlation with all viral activity markers. This result further supports that viral infection enhances TEP production, which in turn stimulates host aggregation [77, 82]. However, a strong positive correlation was only observed between RdRp expression in sinking and POC/PON fluxes (Fig. 6b), suggesting that the impact of viral infection on TEP production—and its effect on POC/PON fluxes—depends on both the host and virus type, with RNA virus infection potentially enhancing particle sinking at the study sites.

**Figure 6.**
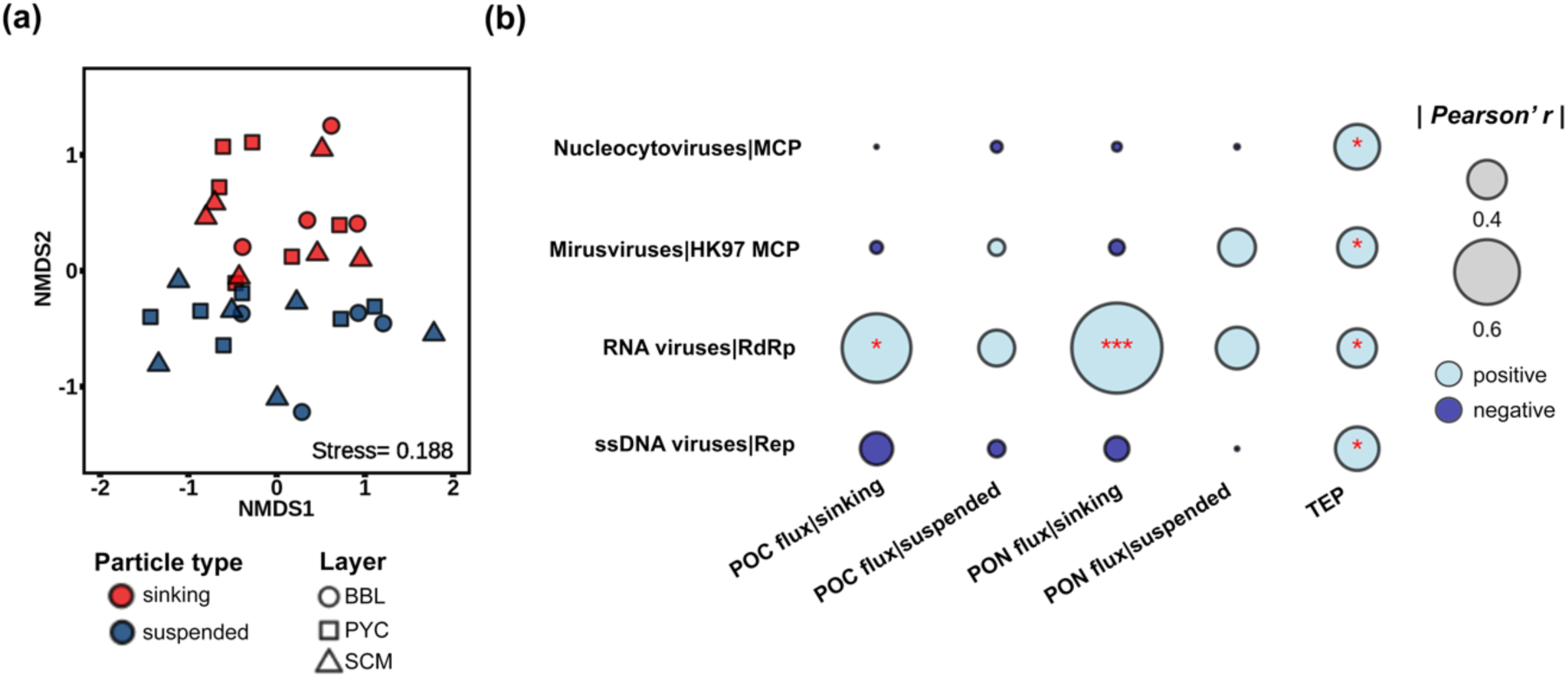
Activity of protistan viruses. (A) nMDS plot showing differences in viral transcript composition between sinking and suspended particles, and across depths, based on transcript expression levels (TPM-normalized within viral transcripts). (B) Pearson’s correlation (r) between viral marker gene expression levels and environmental variables. Abbreviations: POC, particulate organic carbon; PON, particulate organic nitrogen; TEP, transparent exopolymer particles. Asterisks indicate significance levels: * *p*<0.05, ** *p*<0.01, *** *p*<0.001. “POC/PON flux|suspended/sinking” refers to the correlation between POC/PON flux and marker gene expression in corresponding suspended/sinking particles.

In summary, we investigated the role of protists and viruses in the BCP by analyzing differential gene expression between sinking and suspended particles. Our findings revealed significant differences in the metabolic activity of protistan communities and viral activities between these two particle types. Suspended particles exhibited up-regulated DEGs associated with diatoms, haptophytes, and chlorophytes, whereas sinking particles showed up-regulated DEGs related to diatoms, ciliates, and dinoflagellates. These results highlight the selective structuring of protistan communities within the sinking particle microbiome, with viral infection potentially playing a contributing role. Further species-specific examination uncovered substantial differences in transcription profiles between sinking and suspended particles at the species level, indicating niche-dependent cellular metabolic reprogramming. Notably, *M. variabilis* and *T. oceanica* displayed more DEGs in suspended particles, while diatom *T. rotula* exhibited increased gene expression in sinking particles. The presence of both phototrophic and heterotrophic protist activity in sinking particles suggests the export of fresh organic carbon, with heterotrophs facilitating degradation. Moreover, *K. veneficum* and *K. brevis* displayed enhanced gene expression in sinking particles compared to suspended particles, particularly for genes associated with feeding and organic carbon metabolism. This observation implies that dinoflagellate species actively consume organic carbon within sinking particles. Finally, we show that RNA viruses may play a role in promoting particle sinking.

## Acknowledgements

We sincerely thank the captain, officers, and crew of the R/V Shinsei-Maru for their exceptional support and assistance throughout the cruise. This study was funded by JSPS KAKENHI (nos. 19H05667, 22H00384, and 22H02420), JST CREST (JPMJCR23J4), JST FOREST (JPMJFR2070), and World Premier International Research Center Initiative, MEXT, Japan. Computational work was performed at the SuperComputer System, Institute for Chemical Research, Kyoto University.

## Notes

### Competing Interest Statement

The authors have declared no competing interest.

## Reference

1. Volk T, Hoffert MI. Ocean Carbon Pumps: Analysis of Relative Strengths and Efficiencies in Ocean-Driven Atmospheric CO2 Changes. The Carbon Cycle and Atmospheric CO2: Natural Variations Archean to Present. American Geophysical Union (AGU), 1985, 99–110.

2. Korb RE et al. Regional and seasonal differences in microplankton biomass, productivity, and structure across the Scotia Sea: Implications for the export of biogenic carbon. Deep Sea Research Part II: Topical Studies in Oceanography 2012;59–60:67–77. 10.1016/j.dsr2.2011.06.006

3. Worden AZ et al. Rethinking the marine carbon cycle: Factoring in the multifarious lifestyles of microbes. Science 2015;347:1257594. 10.1126/science.1257594

4. Steinberg DK, Landry MR. Zooplankton and the Ocean Carbon Cycle. Annual Review of Marine Science 2017;9:413–444. 10.1146/annurev-marine-010814-015924

5. Smith DC et al. Intense hydrolytic enzyme activity on marine aggregates and implications for rapid particle dissolution. Nature 1992;359:139–142. 10.1038/359139a0

6. Steinberg DK et al. Bacterial vs. zooplankton control of sinking particle flux in the ocean’s twilight zone. Limnology and Oceanography 2008;53:1327–1338. 10.4319/lo.2008.53.4.1327

7. DeLong EF, Franks DG, Alldredge AL. Phylogenetic diversity of aggregate-attached vs. free-living marine bacterial assemblages. Limnology and Oceanography 1993;38:924–934. 10.4319/lo.1993.38.5.0924

8. Lampitt RS et al. Marine snow studies in the Northeast Atlantic Ocean: distribution, composition and role as a food source for migrating plankton. Marine Biology 1993;116:689–702. 10.1007/BF00355486

9. Bidle KD, Fletcher M. Comparison of free-living and particle-associated bacterial communities in the chesapeake bay by stable low-molecular-weight RNA analysis. Applied and Environmental Microbiology 1995;61:944–952. 10.1128/aem.61.3.944-952.1995

10. Amacher J et al. Molecular approach to determine contributions of the protist community to particle flux. Deep Sea Research Part I: Oceanographic Research Papers 2009;56:2206–2215. 10.1016/j.dsr.2009.08.007

11. Ganesh S et al. Metagenomic analysis of size-fractionated picoplankton in a marine oxygen minimum zone. The ISME Journal 2014;8:187–211. 10.1038/ismej.2013.144

12. Duret MT, Lampitt RS, Lam P. Prokaryotic niche partitioning between suspended and sinking marine particles. Environmental Microbiology Reports 2019;11:386–400. 10.1111/1758-2229.12692

13. Gutierrez-Rodriguez A et al. High contribution of Rhizaria (Radiolaria) to vertical export in the California Current Ecosystem revealed by DNA metabarcoding. ISME J 2019;13:964–976. 10.1038/s41396-018-0322-7

14. Yang Q et al. Taxon-specific contributions of microeukaryotes to biological carbon pump in the Oyashio region. ISME Communications 2024;4:ycae136. 10.1093/ismeco/ycae136

15. Poff KE et al. Microbial dynamics of elevated carbon flux in the open ocean’s abyss. Proceedings of the National Academy of Sciences 2021;118:e2018269118. 10.1073/pnas.2018269118

16. Leu AO et al. Diverse Genomic Traits Differentiate Sinking-Particle-Associated versus Free-Living Microbes throughout the Oligotrophic Open Ocean Water Column. mBio 2022;13:e01569–22. 10.1128/mbio.01569-22

17. Buchan A et al. Master recyclers: features and functions of bacteria associated with phytoplankton blooms. Nat Rev Microbiol 2014;12:686–698. 10.1038/nrmicro3326

18. Smith SR, Abbriano RM, Hildebrand M. Comparative analysis of diatom genomes reveals substantial differences in the organization of carbon partitioning pathways. Algal Research 2012;1:2–16. 10.1016/j.algal.2012.04.003

19. Jacobson DM, Anderson DM. Widespread Phagocytosis of Ciliates and Other Protists by Marine Mixotrophic and Heterotrophic Thecate Dinoflagellates. Journal of Phycology 1996;32:279–285. 10.1111/j.0022-3646.1996.00279.x

20. Labarre A et al. Expression of genes involved in phagocytosis in uncultured heterotrophic flagellates. Limnology and Oceanography 2020;65:S149–S160. 10.1002/lno.11379

21. Stoecker DK et al. Mixotrophy in the Marine Plankton. Ann Rev Mar Sci 2017;9:311–335. 10.1146/annurev-marine-010816-060617

22. Jeong HJ et al. Growth, feeding and ecological roles of the mixotrophic and heterotrophic dinoflagellates in marine planktonic food webs. Ocean Sci J 2010;45:65–91. 10.1007/s12601-010-0007-2

23. Ismail WM, Ye Y, Tang H. Gene finding in metatranscriptomic sequences. BMC Bioinformatics 2014;15:S8. 10.1186/1471-2105-15-S9-S8

24. Yamada Y et al. Functions of extracellular polymeric substances in partitioning suspended and sinking particles in the upper oceans of two open ocean systems. Limnology and Oceanography 2024;69:1101–1114. 10.1002/lno.12554

25. Chen S et al. fastp: an ultra-fast all-in-one FASTQ preprocessor. Bioinformatics 2018;34:i884–i890. 10.1093/bioinformatics/bty560

26. Xu H et al. FastUniq: A Fast De Novo Duplicates Removal Tool for Paired Short Reads. PLOS ONE 2012;7:e52249. 10.1371/journal.pone.0052249

27. Kopylova E, Noé L, Touzet H. SortMeRNA: fast and accurate filtering of ribosomal RNAs in metatranscriptomic data. Bioinformatics 2012;28:3211–3217. 10.1093/bioinformatics/bts611

28. Haas BJ et al. De novo transcript sequence reconstruction from RNA-seq using the Trinity platform for reference generation and analysis. Nat Protoc 2013;8:1494–1512. 10.1038/nprot.2013.084

29. Rice P, Longden I, Bleasby A. EMBOSS: The European Molecular Biology Open Software Suite. Trends in Genetics 2000;16:276–277. 10.1016/S0168-9525(00)02024-2

30. Steinegger M, Söding J. MMseqs2 enables sensitive protein sequence searching for the analysis of massive data sets. Nat Biotechnol 2017;35:1026– 1028. 10.1038/nbt.3988

31. Patro R et al. Salmon provides fast and bias-aware quantification of transcript expression. Nat Methods 2017;14:417–419. 10.1038/nmeth.4197

32. Buchfink B, Xie C, Huson DH. Fast and sensitive protein alignment using DIAMOND. Nat Methods 2015;12:59–60. 10.1038/nmeth.3176

33. Groussman RD et al. MarFERReT, an open-source, version-controlled reference library of marine microbial eukaryote functional genes. Sci Data 2023;10:926. 10.1038/s41597-023-02842-4

34. O’Leary NA et al. Reference sequence (RefSeq) database at NCBI: current status, taxonomic expansion, and functional annotation. Nucleic Acids Res 2016;44:D733–D745. 10.1093/nar/gkv1189

35. Zayed AA et al. Cryptic and abundant marine viruses at the evolutionary origins of Earth’s RNA virome. Science 2022;376:156–162. 10.1126/science.abm5847

36. Gaïa M et al. Mirusviruses link herpesviruses to giant viruses. Nature 2023;616:783–789. 10.1038/s41586-023-05962-4

37. Kanehisa M. KEGG: Kyoto Encyclopedia of Genes and Genomes. Nucleic Acids Research 2000;28:27–30. 10.1093/nar/28.1.27

38. Huerta-Cepas J et al. eggNOG 5.0: a hierarchical, functionally and phylogenetically annotated orthology resource based on 5090 organisms and 2502 viruses. Nucleic Acids Research 2019;47:D309–D314. 10.1093/nar/gky1085

39. Hu SK et al. Shifting metabolic priorities among key protistan taxa within and below the euphotic zone. Environmental Microbiology 2018;20:2865–2879. 10.1111/1462-2920.14259

40. Robinson MD, Oshlack A. A scaling normalization method for differential expression analysis of RNA-seq data. Genome Biol 2010;11:R25. 10.1186/gb-2010-11-3-r25

41. Dixon P. VEGAN, a package of R functions for community ecology. Journal of Vegetation Science 2003;14:927–930. 10.1111/j.1654-1103.2003.tb02228.x

42. Robinson MD, McCarthy DJ, Smyth GK. edgeR : a Bioconductor package for differential expression analysis of digital gene expression data. Bioinformatics 2010;26:139–140. 10.1093/bioinformatics/btp616

43. Benjamini Y, Hochberg Y. Controlling the False Discovery Rate: A Practical and Powerful Approach to Multiple Testing. Journal of the Royal Statistical Society: Series B (Methodological) 1995;57:289–300. 10.1111/j.2517-6161.1995.tb02031.x

44. Camargo AP et al. Identification of mobile genetic elements with geNomad. Nat Biotechnol 2024;42:1303–1312. 10.1038/s41587-023-01953-y

45. Weitz JS, Levin SA. Size and scaling of predator–prey dynamics. Ecology Letters 2006;9:548–557. 10.1111/j.1461-0248.2006.00900.x

46. Tsai C-H, Hsieh C, Nakazawa T. Predator–prey mass ratio revisited: does preference of relative prey body size depend on individual predator size? Functional Ecology 2016;30:1979–1987. 10.1111/1365-2435.12680

47. Deretic V. Autophagosome and Phagosome. In: Deretic V (ed.), Autophagosome and Phagosome. Totowa, NJ: Humana Press, 2008, 1–10.

48. Lawrence RE, Zoncu R. The lysosome as a cellular centre for signalling, metabolism and quality control. Nat Cell Biol 2019;21:133–142. 10.1038/s41556-018-0244-7

49. Rubin ET et al. Transcriptomic Response to Feeding and Starvation in a Herbivorous Dinoflagellate. Front Mar Sci 2019;6. 10.3389/fmars.2019.00246

50. Shanks A, Walters K. Feeding by a heterotrophic dinoflagellate (Noctiluca scintillans) in marine snow. Limnology and Oceanography 1996;41:177–181. 10.4319/lo.1996.41.1.0177

51. Scott M et al. Interdependence of Cell Growth and Gene Expression: Origins and Consequences. Science 2010;330:1099–1102. 10.1126/science.1192588

52. Smetacek V, Assmy P, Henjes J. The role of grazing in structuring Southern Ocean pelagic ecosystems and biogeochemical cycles. Antarctic Science 2004;16:541–558. 10.1017/S0954102004002317

53. Bock NA et al. Experimental identification and in silico prediction of bacterivory in green algae. The ISME Journal 2021;15:1987–2000. 10.1038/s41396-021-00899-w

54. Pang M, Liu K, Liu H. Evidence for mixotrophy in pico-chlorophytes from a new Picochlorum (Trebouxiophyceae) strain. Journal of Phycology 2022;58:80–91. 10.1111/jpy.13218

55. Artolozaga I et al. Grazing rates of bacterivorous protists inhabiting diverse marine planktonic microenvironments. Limnology and Oceanography 2002;47:142–150. 10.4319/lo.2002.47.1.0142

56. Bar-Zeev E et al. Programmed cell death in the marine cyanobacterium *Trichodesmium* mediates carbon and nitrogen export. The ISME Journal 2013;7:2340–2348. 10.1038/ismej.2013.121

57. Bidle KD. Programmed Cell Death in Unicellular Phytoplankton. Current Biology 2016;26:R594–R607. 10.1016/j.cub.2016.05.056

58. Agusti S et al. Ubiquitous healthy diatoms in the deep sea confirm deep carbon injection by the biological pump. Nat Commun 2015;6:7608. 10.1038/ncomms8608

59. Vasiljeva O et al. Emerging roles of cysteine cathepsins in disease and their potential as drug targets. *Curr Pharm Des* 2007;13:387–403. 10.2174/138161207780162962

60. Turk V et al. Cysteine cathepsins: from structure, function and regulation to new frontiers. Biochim Biophys Acta 2012;1824:68–88. 10.1016/j.bbapap.2011.10.002

61. Doherty GJ, McMahon HT. Mechanisms of Endocytosis. Annual Review of Biochemistry 2009;78:857–902. 10.1146/annurev.biochem.78.081307.110540

62. Kumari S, Mg S, Mayor S. Endocytosis unplugged: multiple ways to enter the cell. Cell Res 2010;20:256–275. 10.1038/cr.2010.19

63. Kazamia E et al. Endocytosis-mediated siderophore uptake as a strategy for Fe acquisition in diatoms. Science Advances 2018;4:eaar4536. 10.1126/sciadv.aar4536

64. McQuaid JB et al. Carbonate-sensitive phytotransferrin controls high-affinity iron uptake in diatoms. Nature 2018;555:534–537. 10.1038/nature25982

65. Lampe RH et al. Molecular Mechanisms for Iron Uptake and Homeostasis in Marine Eukaryotic Phytoplankton. Annual Review of Microbiology 2024;78:213–232. 10.1146/annurev-micro-041222-023252

66. Esteban GF, Fenchel T, Finlay BJ. Mixotrophy in Ciliates. Protist 2010;161:621–641. 10.1016/j.protis.2010.08.002

67. Granéli E, Johansson N. Effects of the toxic haptophyte Prymnesium parvum on the survival and feeding of a ciliate: the influence of different nutrient conditions. Marine Ecology Progress Series 2003;254:49–56. 10.3354/meps254049

68. Aberle N, Lengfellner K, Sommer U. Spring bloom succession, grazing impact and herbivore selectivity of ciliate communities in response to winter warming. Oecologia 2007;150:668–681. 10.1007/s00442-006-0540-y

69. Sherr EB, Sherr BF. Heterotrophic dinoflagellates: a significant component of microzooplankton biomass and major grazers of diatoms in the sea. Marine Ecology Progress Series 2007;352:187–197. 10.3354/meps07161

70. Hinder SL et al. Changes in marine dinoflagellate and diatom abundance under climate change. Nature Clim Change 2012;2:271–275. 10.1038/nclimate1388

71. Li Y, Guo X-H, Lundholm N. Molecular Phylogeny and Taxonomy of the Genus Minidiscus (Bacillariophyceae), with Description of Mediolabrus gen. nov. Journal of Phycology 2020;56:1443–1456. 10.1111/jpy.13038

72. Lommer M et al. Genome and low-iron response of an oceanic diatom adapted to chronic iron limitation. Genome Biology 2012;13:R66. 10.1186/gb-2012-13-7-r66

73. Wyatt SN, McNabb, Brandon J., and Varela DE. Morphological and physiological responses of the cosmopolitan marine diatom Thalassiosira rotula to acidification. Diatom Research 2024;39:61–74. 10.1080/0269249X.2024.2369049

74. Tréguer P et al. Influence of diatom diversity on the ocean biological carbon pump. Nature Geosci 2018;11:27–37. 10.1038/s41561-017-0028-x

75. Dl M, M L, Pj H. Effects of iron and nitrogen source on the sinking rate, physiology and metal composition of an oceanic diatom from the subarctic Pacific. Marine Ecology Progress Series 1996;132:215–227. 10.3354/meps132215

76. Pahlow M, Riebesell U, Wolf-Gladrow DA. Impact of cell shape and chain formation on nutrient acquisition by marine diatoms. Limnology and Oceanography 1997;42:1660–1672. 10.4319/lo.1997.42.8.1660

77. Yamada Y et al. Aggregate Formation During the Viral Lysis of a Marine Diatom. Front Mar Sci 2018;5. 10.3389/fmars.2018.00167

78. Ressurreição M et al. A role for p38 MAPK in the regulation of ciliary motion in a eukaryote. BMC Cell Biology 2011;12:6. 10.1186/1471-2121-12-6

79. Gowans GJ et al. AMP Is a True Physiological Regulator of AMP-Activated Protein Kinase by Both Allosteric Activation and Enhancing Net Phosphorylation. Cell Metabolism 2013;18:556–566. 10.1016/j.cmet.2013.08.019

80. Calbet A et al. Intraspecific variability in *Karlodinium veneficum*: Growth rates, mixotrophy, and lipid composition. Harmful Algae 2011;10:654–667. 10.1016/j.hal.2011.05.001

81. Ahn SH, Glibert PM. Temperature-Dependent Mixotrophy in Natural Populations of the Toxic Dinoflagellate Karenia brevis. Water 2024;16:1555. 10.3390/w16111555

82. Lønborg C, Middelboe M, Brussaard CPD. Viral lysis of Micromonas pusilla: impacts on dissolved organic matter production and composition. Biogeochemistry 2013;116:231–240. 10.1007/s10533-013-9853-1

