## Supplementary Figs. S1-S12 for "Metabolic reprogramming of protists within the microbiome of sinking particles"

**Running title:** Metabolic Reprogramming in Sinking Particle Microbiomes

*Corresponding authors:

This supplementary file includes:

Supplementary Figs. S1-S12

Legends of Supplementary Tables S1-S4

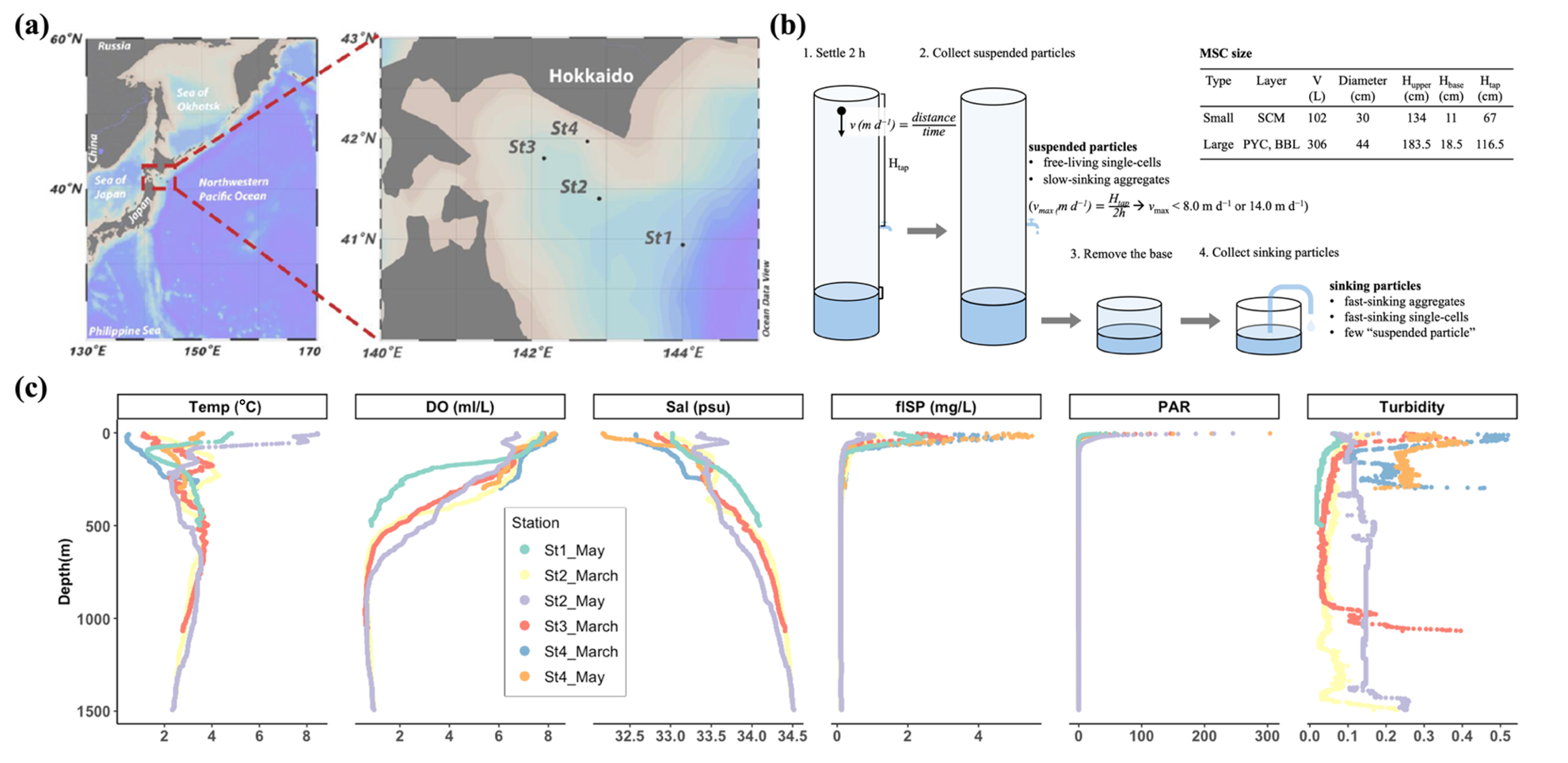

**Figure S1** Overview of the Sampling Strategy. (A) Sampling site locations displayed using Ocean Data View. (B) Diagram illustrating the deployment of Marine Snow Catchers (MSC). Two types of MSCs were utilized: a smaller MSC for the SCM layer and a larger MSC for the PYC and BBL layers, with specifications provided in the accompanying table. (C) Profiles of water column properties obtained via CTD profiling, including temperature (Temp), dissolved oxygen (DO), salinity (Sal), chlorophyll fluorescence (flSP), photosynthetically active radiation (PAR), and turbidity measurements.

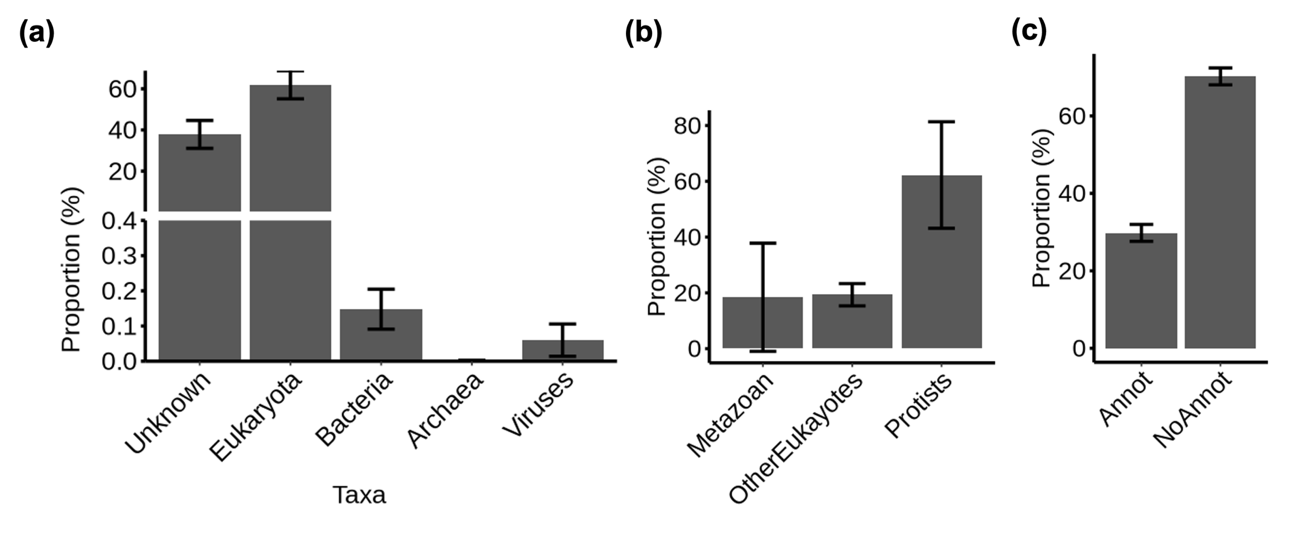

**Figure S2** Taxonomic and functional assignment of transcripts. (A) Proportion of taxonomic assignments for all transcripts. (B) Proportion of taxonomic assignments for all eukaryotic transcripts. (C) Proportion of KO hits for all protistan transcripts.

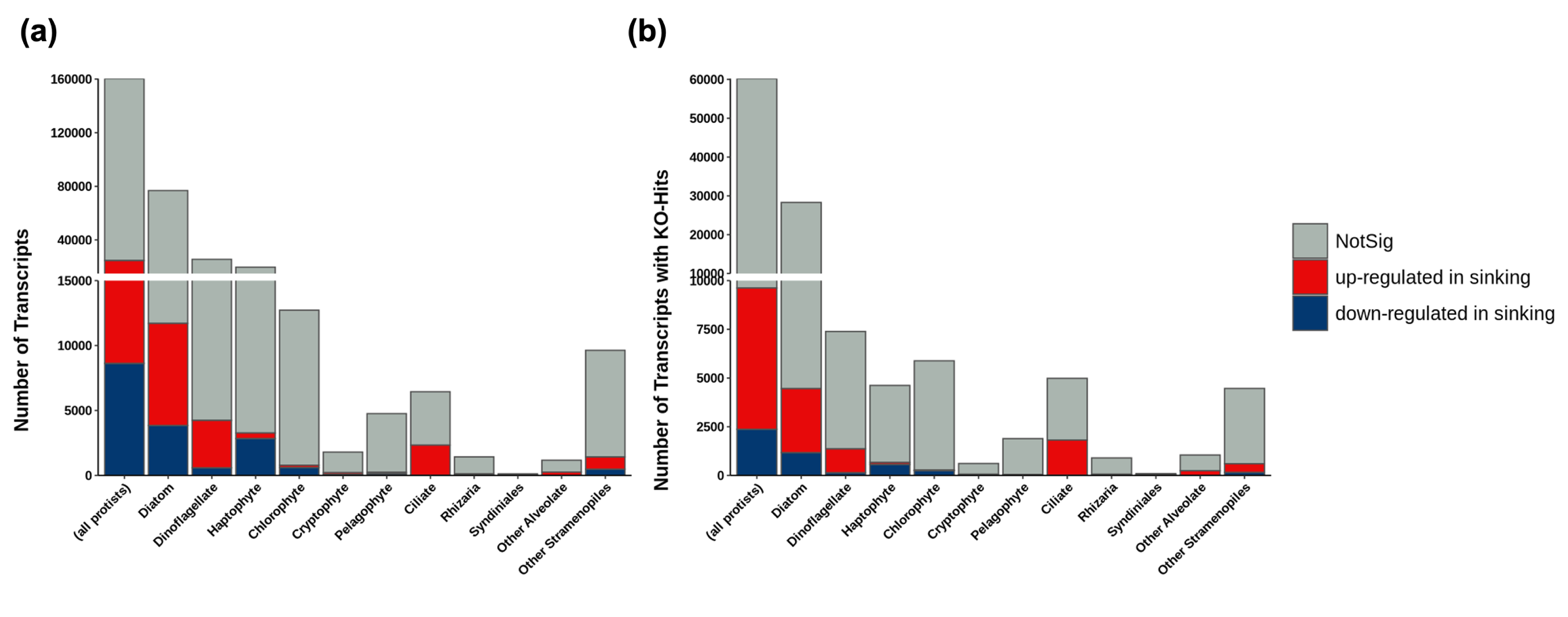

**Figure S3** Counts of differentially expressed genes between sinking and suspended particles across taxa. (A) All transcripts identified by edgeR. (B) Only KO-hits transcripts identified by edgeR.

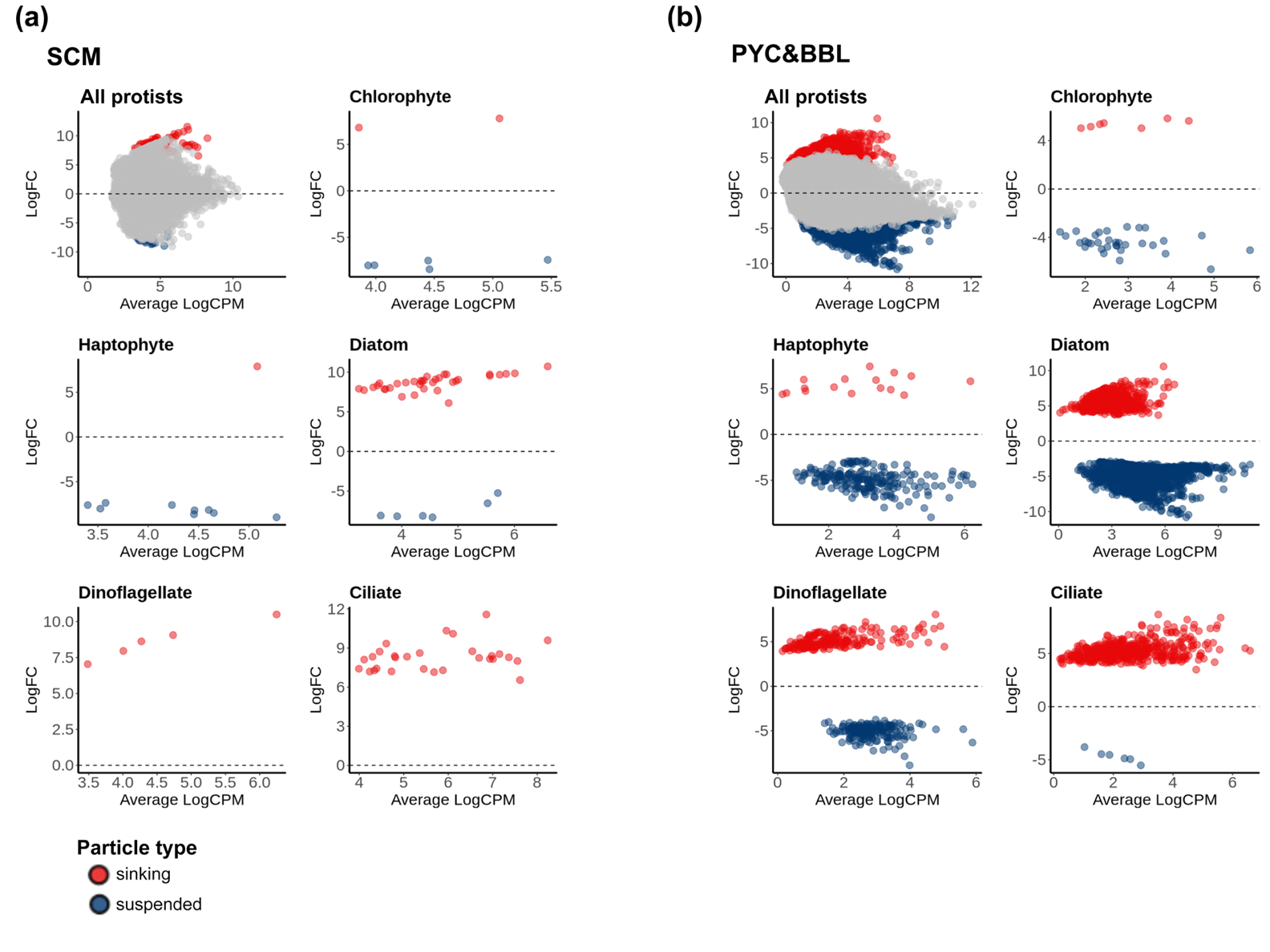

**Figure S4** MA plots show differential gene expression between sinking and suspended particles for all protists and by major taxa (chlorophyte, haptophyte, diatom, dinoflagellate, and ciliate). The x-axis represents the mean log-transformed counts per million (CPM) from TMM-normalized libraries (only KO-annotated transcripts included). (A) Samples from the SCM layer. (B) Samples from the PYC and BBL layers.

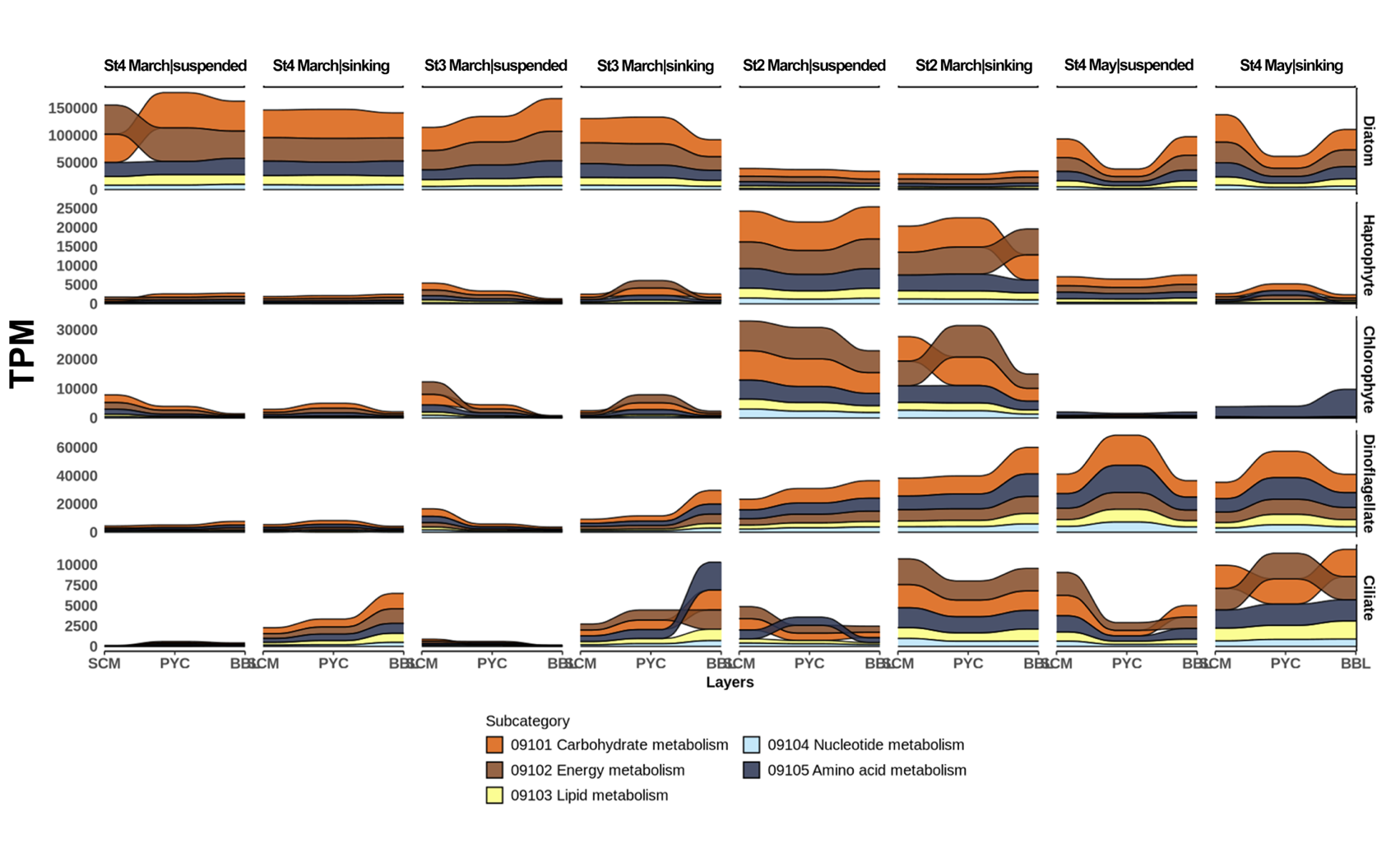

**Figure S5** Vertical changes in the expression of major metabolic groups across taxa in sinking and suspended particles. Transcript expression was normalized to TPM. The metabolic groups, including carbohydrate metabolism, energy metabolism, lipid metabolism, nucleotide metabolism, and amino acid metabolism, were compiled from the KEGG database. Stations without a BBL sample (St1 and St2 in May) were excluded from the analysis.

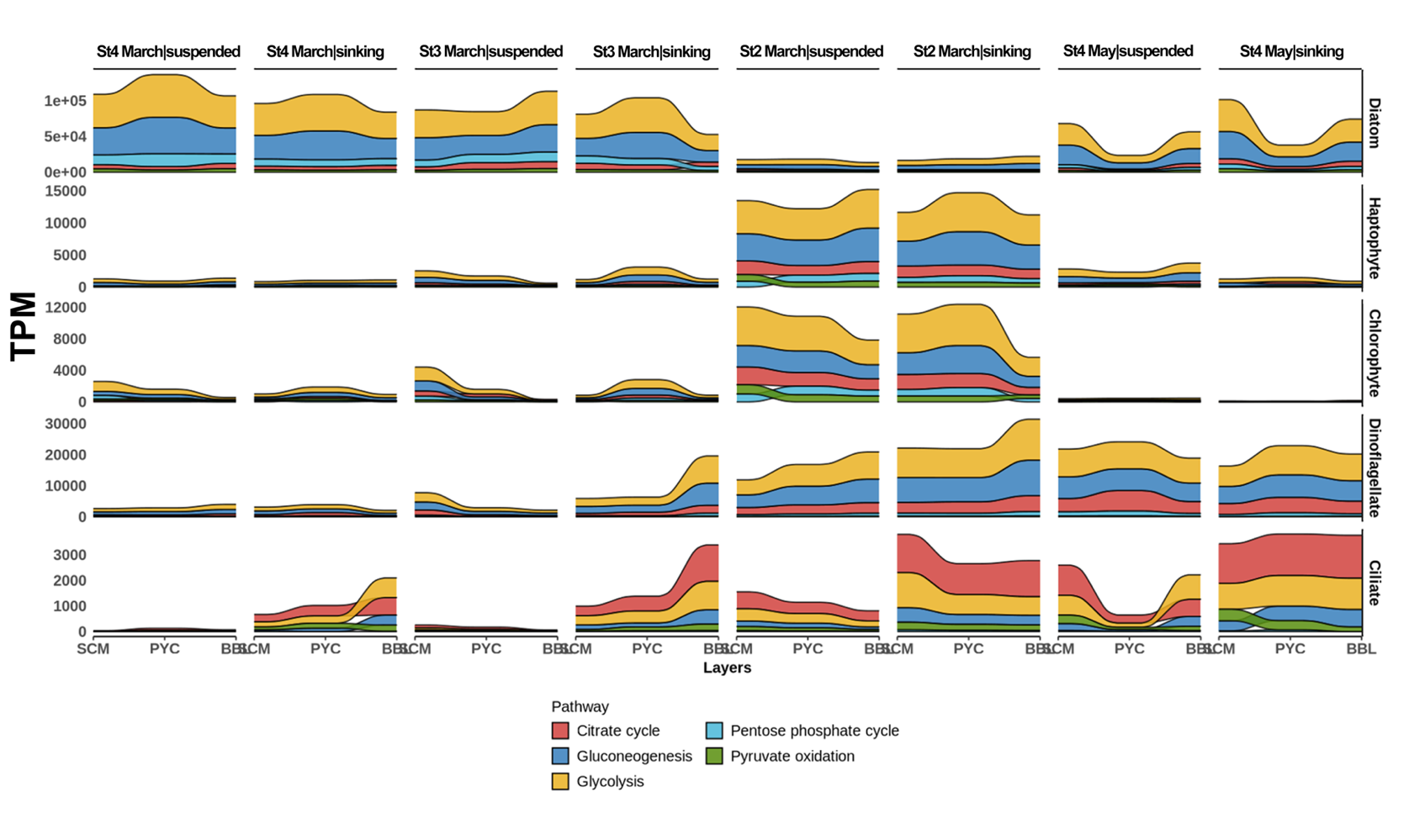

**Figure S6** Vertical changes in the expression of major carbohydrate metabolism pathways across taxa in sinking and suspended particles. Transcript expression was normalized to TPM. The carbohydrate metabolism pathways, including the citrate cycle, gluconeogenesis, glycolysis, pentose phosphate pathway, and pyruvate oxidation, were compiled from the KEGG database. Stations without a BBL sample (St1 and St2 in May) were excluded from the analysis.

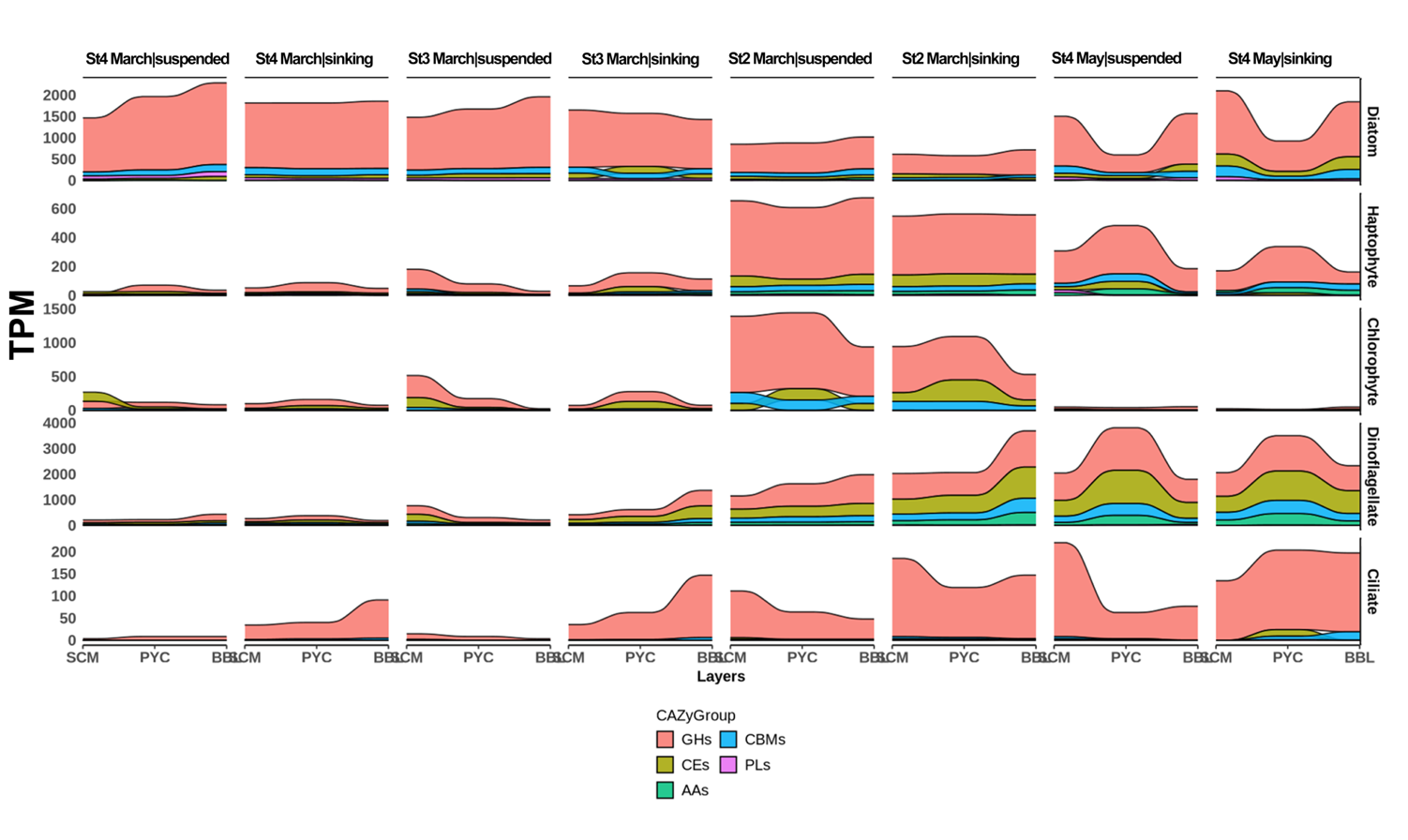

**Figure S7** Vertical changes in the expression of enzyme families related to carbohydrate degradation based on CAZymes across taxa in sinking and suspended particles. Transcript expression was normalized to TPM. The putative carbohydrate-degrading capacity, based on CAZymes, includes glycoside hydrolases (GHs), carbohydrate esterases (CEs), polysaccharide lyases (PLs), carbohydrate-binding modules (CBMs), and auxiliary activities (AAs). Stations without a BBL sample (St1 and St2 in May) were excluded from the analysis.

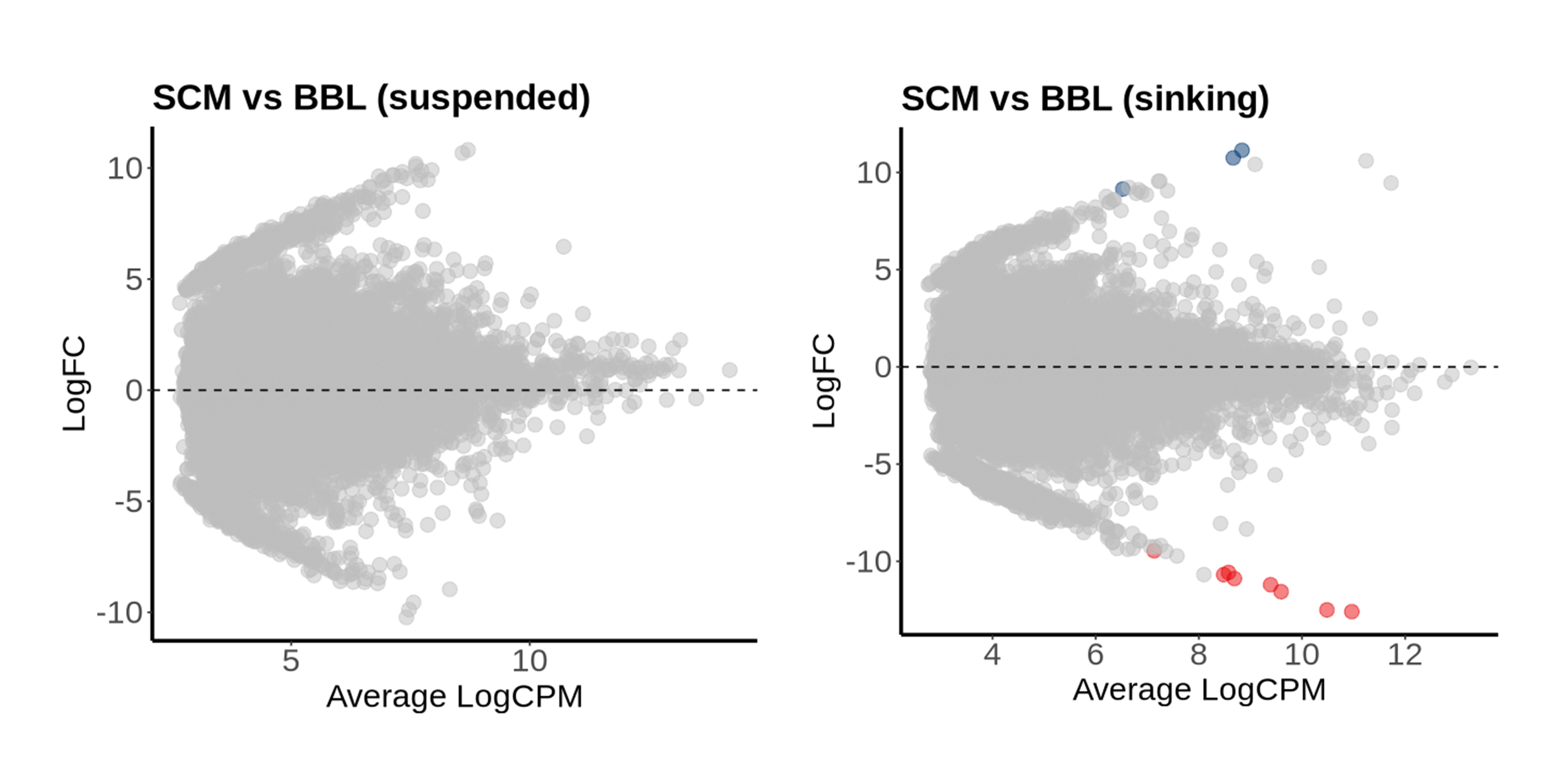

**Figure S8** MA plots showing differential gene expression of phototrophy- and heterotrophy-related genes between SCM and BBL for all protists. The x-axis represents the mean log-transformed CPM from TMM-normalized libraries.

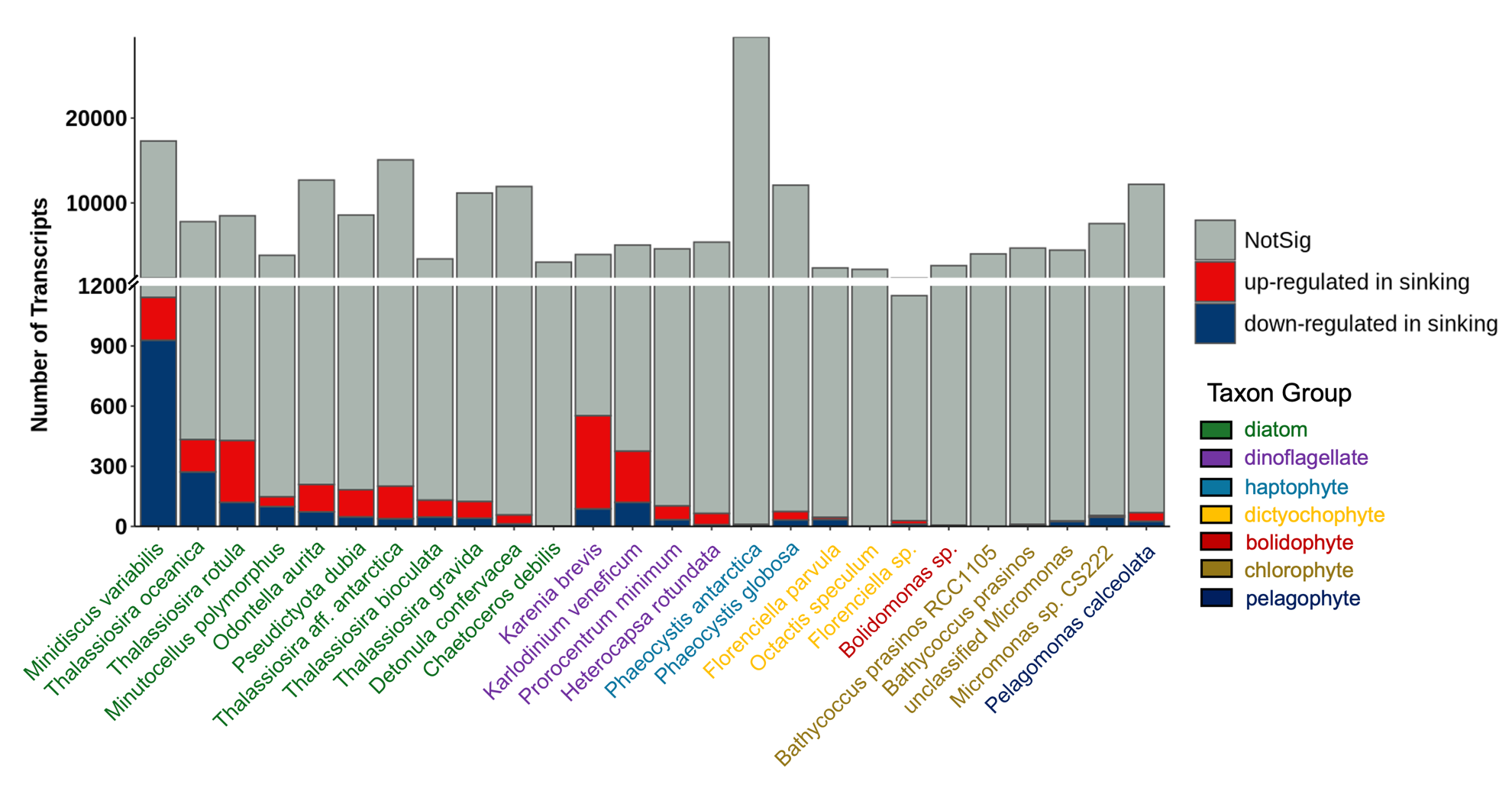

**Figure S9** Counts of differentially expressed genes between sinking and suspended particles across species-level transcript bins.

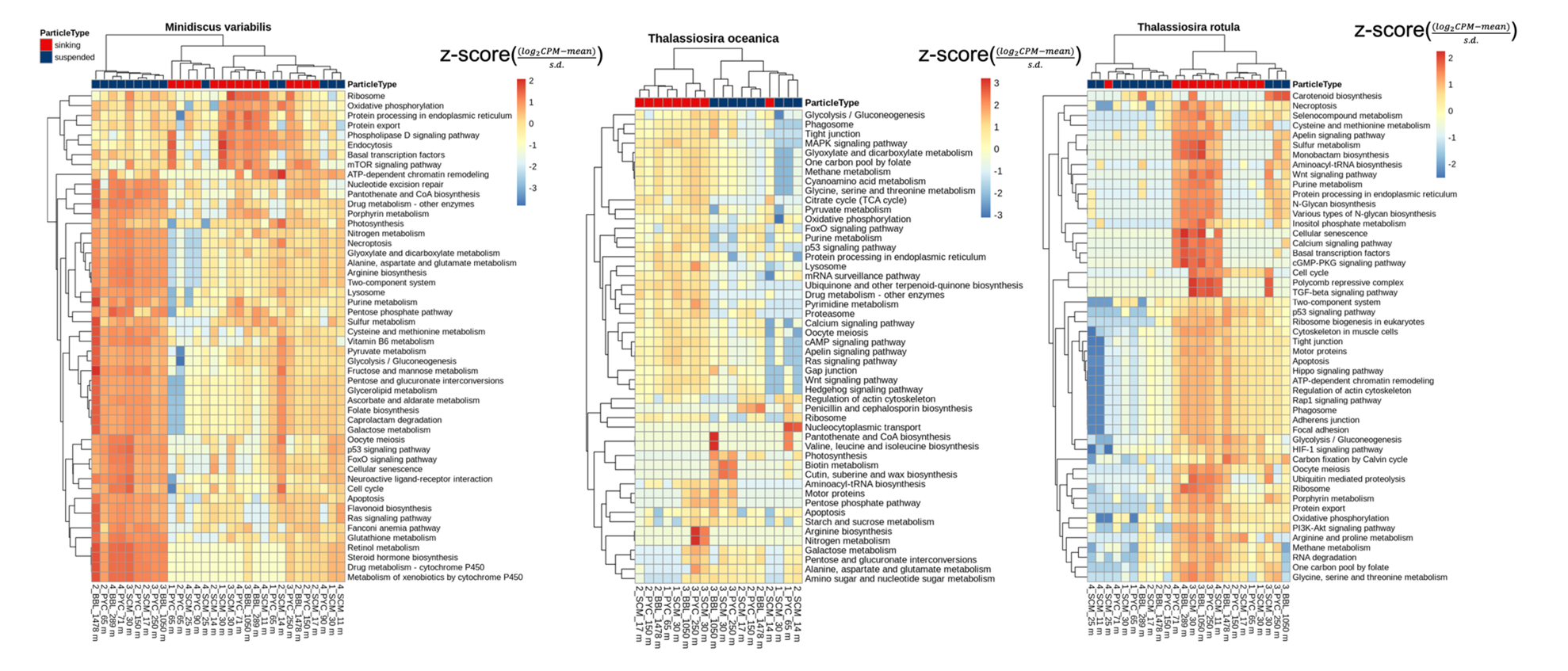

**Figure S10** Heatmap showing the expression of top 50 KEGG pathways associated with differentially expressed transcripts with KO hits identified between sinking and suspended particles in diatom species. Differential expression is represented as the z-score ((log_2_CPM - mean) / s.d.).

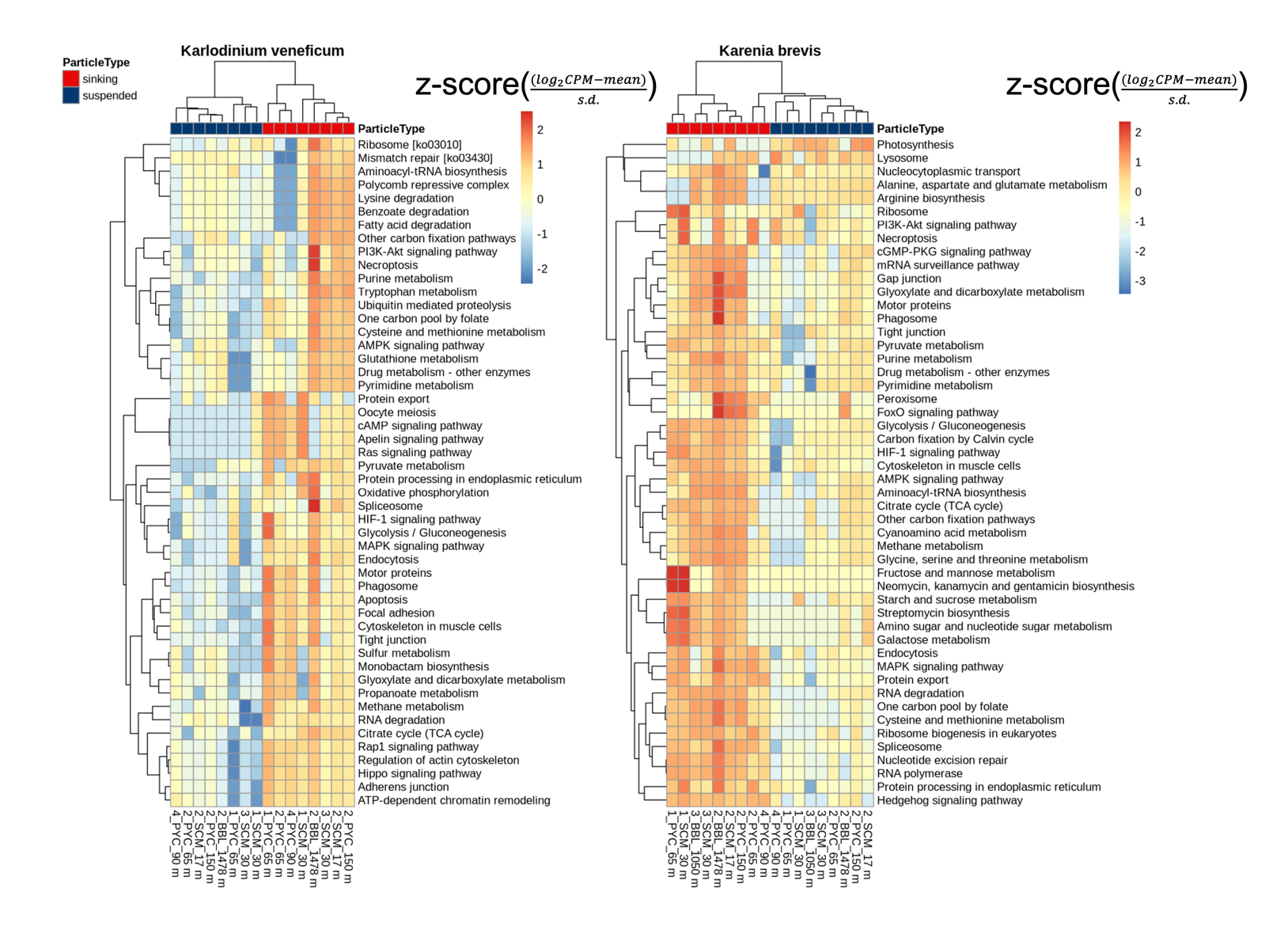

**Figure S11** Heatmap showing the expression of top 50 KEGG pathways associated with differentially expressed transcripts with KO hits identified between sinking and suspended particles in dinoflagellate species. Differential expression is represented as the z-score ((log_2_CPM - mean) / s.d.).

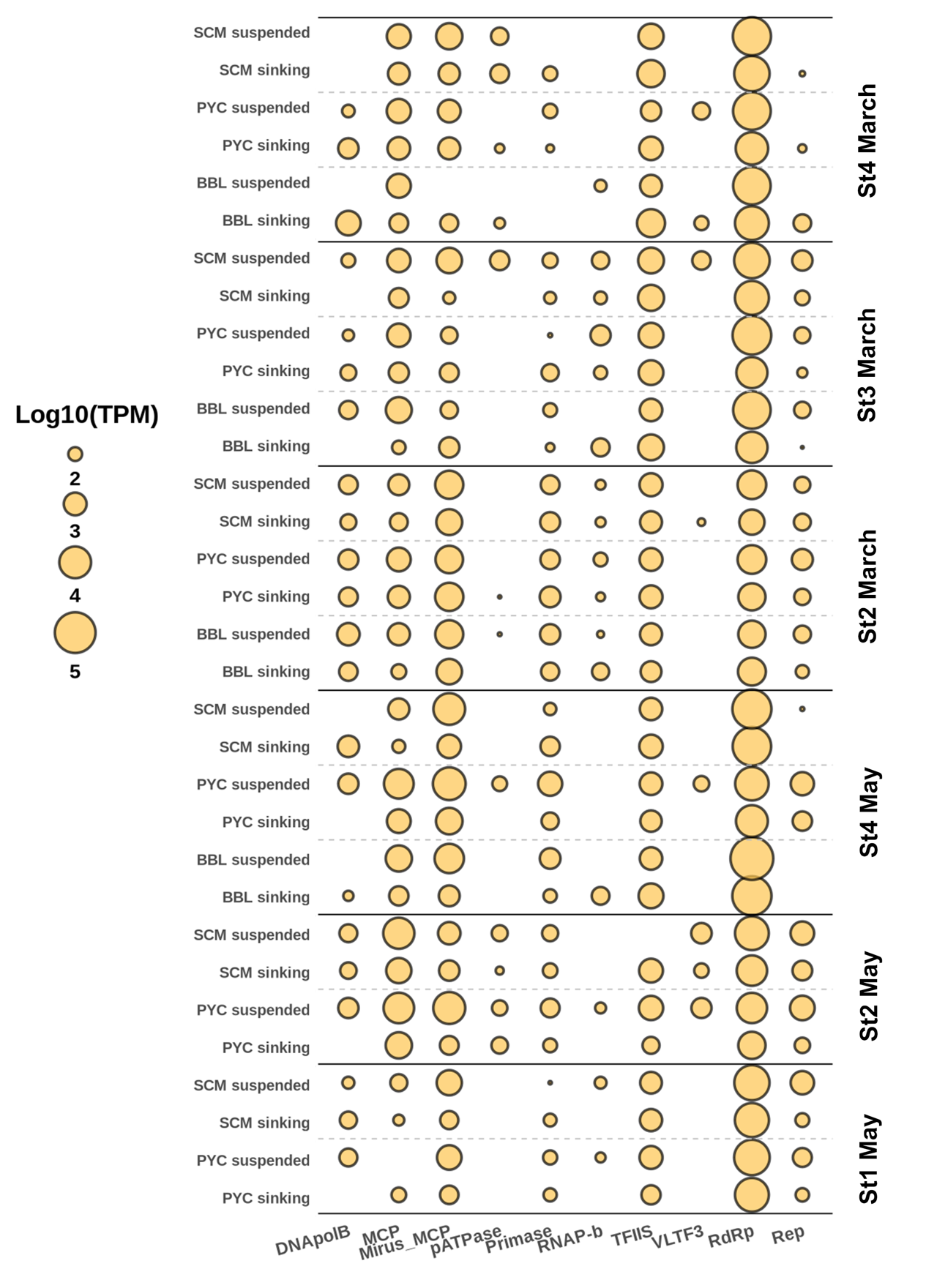

**Figure S12** Heatmap of expression level (TPM-normalized within viral transcripts) of viral marker genes across samples.

**Table S1** Detail sampling information and summary statistics of the Metatranscriptome sequencing data processing. MSC type refers to the MSC size used for sampling; small MSC with 100 L upper part and 20 L base part; giant MSC with 300 L upper part and 70L base part.

**Table S2** The KO list of biomarker gene groups for phototrophy and heterotrophy. The corresponding KEGG KO terms, gene names, biomarker categories, and relevant references for each gene are provided.

**Table S3** Sources of HMM profiles for marker/core genes of giant viruses and RNA viruses.

**Table S4** PERMANOVA results.
